# Trade-off between branching and polarity controls decision-making during cell migration

**DOI:** 10.1101/2024.03.29.585638

**Authors:** Jiayi Liu, Javier Boix-Campos, Jonathan E. Ron, Johan M. Kux, Magdalena E.M. Oremek, Adriano G. Rossi, Nir S. Gov, Pablo J. Sáez

## Abstract

Migrating cells often face microenvironmental constraints that force them to extend multiple, often highly dynamic, protrusions, that compete to choose the new direction. However, the analysis of how cells coordinate shape dynamics during this directional decision-making process has been restricted to single junctions. Here, we present a theoretical model and the corresponding experimental proof of concept using *in vivo* and *in vitro* live-cell microscopy and a neuronal network-based image analysis pipeline, to explore the shape and migration dynamics of highly bifurcated cells during spontaneous random migration. We found that macrophages and endothelial cells display different migration regimes in a hexagonal adhesive network, despite sharing a mesenchymal migratory strategy. Macrophages moved faster and presented larger changes in cell length in comparison to endothelial cells. The theoretical model describes the behavior of both cells during directional decision-making, and it reveals a trade-off between exploration for directional cues and long-range migration efficiency, showing the fine tune regulation of shape dynamics in complex geometries.

**Teaser:** Highly branched cells require precise control of their shape dynamics to ensure microenvironment exploration while keeping their motility.

## Introduction

Cells migrate within tissues during physiological and pathological processes, such as embryo development, angiogenesis, tissue growth and repair, immune response, and cancer propagation [1–5]. Due to the constraints of the microenvironment, motile cells often need to navigate tissues and organs avoiding the surrounding obstacles [6, 7]. Cell migration in complex geometries has been studied in different *in vitro* models using different geometries and types of confinement [8–17]. Once migrating cells encounter junctions, they generate protrusions along alternate directions [18], and eventually select a new direction in a process named directional decision-making [4, 17, 19]. This response has been observed in different levels of complexity and in different cellular models, such as endothelial[20], immune [21, 22] and cancer [2] cells. However, how highly branched cells can efficiently migrate through complex geometries is still poorly understood.

We have recently shown how cellular shape and cytoskeleton dynamics control the migrating behavior of cells facing a single symmetric Y-junction [17]. Here, we generalize and extend our theoretical model [17] to describe cells that simultaneously span multiple junctions, as frequently occurs inside tissues where similar geometries are encountered. Indeed, during angiogenesis [1], and immune response [23], both endothelial cells and leukocytes are required to migrate inside and over blood vessels, which often bifurcate and force cellular branching [24]. In some cases the junctions offer paths with a bias (i.e. dead end, chemokine, etc), but in some others there is no bias at the junction [17]. Interestingly, unlike other leukocytes, macrophages exhibit a mesenchymal migratory strategy that is also observed in endothelial cells [1, 25], and which is highly dependent on adhesion [23, 26]. Thus, we used these two cell types as models to study branching dynamics during stick-slip migration.

Here, we combined live-cell imaging with micropatterning and advanced image analysis based on a neuronal network to monitor the spontaneous migratory behaviour and shape dynamics of bone marrow-derived macrophages, and human umbilical vein endothelial cells (HUVEC) on adhesive hexagonal networks. We found that both macrophages and HUVEC effectively resolve high levels of branching to continue their migration, although macrophages were more efficient in this process. In addition, we present a coarse-grained theoretical model to describe how highly branched cells form competing arms, and how they are able to maintain their migratory capacity. The model predicts the behavior of both cells during directional decision-making while facing multiple junctions, despite the differences in their migratory behaviour. Our model revealed an important trade-off that affects the optimal amount of cellular branching: a larger number of branches is needed for local sensing of directional cues within the complex microenvironment, but it also decreases the efficiency of the long-range cellular migration.

## The model

To enunciate our model, we considered that some tissues cells have a patterned topology that induces regular branching in migrating cells. Such example is the case of macrophages migrating *in vivo* in the basal layer of keratinocytes in the zebrafish tailfin (Fig. 1A, Movie S1). These keratinocytes have a polygonal shape that often resembles an hexagon. Analysis of the cellular branching shows highly dynamic protrusions, that are coordinated in order to allow proper macrophage tissue surveillance, while preserving migratory capacity (Fig. 1B-C, Movie S2).

**Fig. 1:**
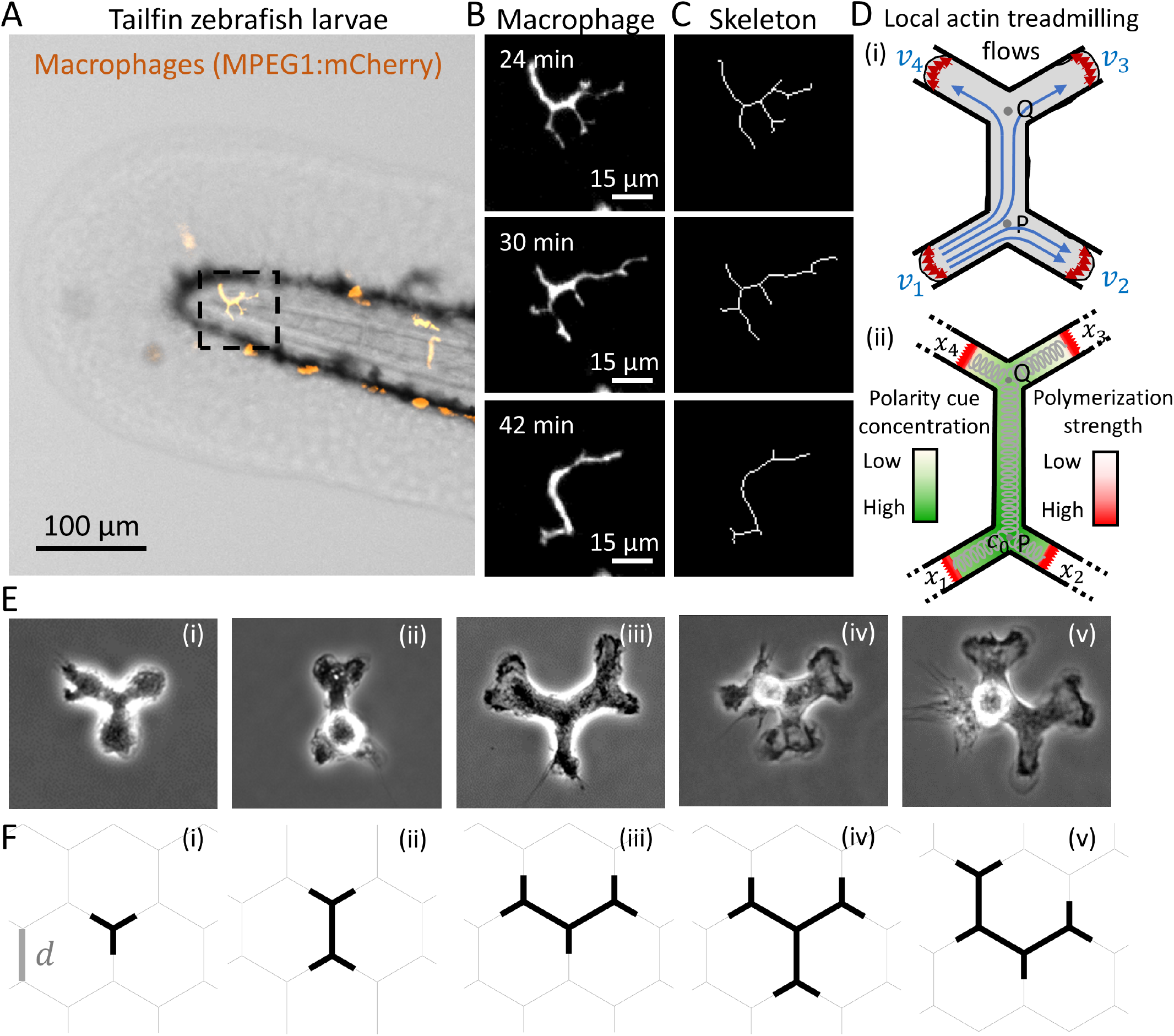
Experimental and theoretical models to study multiple junctions. (A) Image of the tailfin of zebrafish larvae expressing a macrophage marker (MPEG1:mCherry, orange, Movie S1). Dashed square: region of the migrating macrophage shown in (B). (B) Fluorescent images of a macrophage migrating *in vivo* (Movie S2). (C) Skeletonization of the macrophage in B shows the multiple branches (Movie S2). (D) (i) Scheme showing the local actin treadmilling flows at each edge of the arms, and the symmetrical split of the local actin flow that emanates from arm 1 at each junction. (ii) Example of the concentration field of the polarity cue that is affected by the advection flows. Cell elasticity is denoted by the grey springs. (E) Macrophages spanning multiple junctions in adhesive micropatterns. (i)-(v) correspond to the shapes F(i)-(v) respectively. (F) Shape of the cells that are spanning different number of junctions. (i) One, (ii) two, and (iii) three junctions. (iv-v) Two possible cell shapes for cells spanning four junctions.

Our model is based on a cell composed of one-dimensional sections, which spontaneously self-polarize due to the coupling between the local actin polymerization activity at the leading edge of the protrusions (Fig. 1D(i)) [17, 27]. This coupling is mediated by the advection of an inhibitor of actin polymerization (termed “polarity cue”) by the net actin retrograde flow (Fig. 1D(ii) and Fig. S-1). The cell polarizes when one of the tips has the largest actin polymerization activity, which is maintained by the actin flow that advects the inhibitor to the remaining protrusions, where actin polymerization is inhibited.

We extend here our previous model for cells moving over a symmetric Y-shaped junction [17] to describe cells moving on a symmetric hexagonal network of junctions where the cells span up to four junctions simultaneously (Fig. 1E-F and Fig. S-1). The model could also be extended to describe cells spanning a larger number of junctions.

The polymerization of actin at the cellular tips is converted into protrusive forces, which compete with elastic and friction forces. This force balance drives the extension/retraction of the cellular protrusions, and causes the cell to migrate. For simplicity, we assume that all the forces which act on each cellular protrusion are localized at their tips (frontal edges). The global elasticity of the cell is treated as an elastic spring (Fig. 1D(ii)).

For a cell with *N* (*N ≥* 3) arms, i.e., spanning across *N ™* 2 junctions (Fig. 1F), the dynamics of the arm *i* are described by three variables [17]: 1) its length *x*_*i*_, 2) the fraction of active slip-bonds adhesion *n*_*i*_ at the leading edge, and 3) the local actin treadmilling flow velocity *v*_*i*_ at the leading edge (Fig. 1D(i)). The dynamic equations for these variables are given by

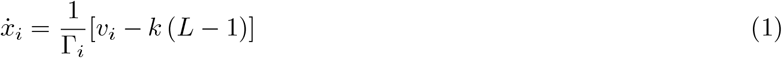

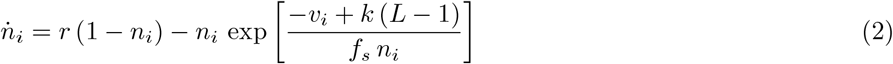

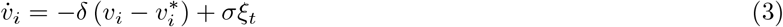

In Eq.1, Γ_*i*_ is a non-constant friction coefficient that depends on the direction of motion of the arm’s leading edge, given by

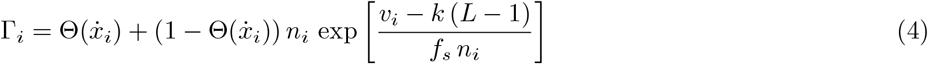

where Θ is a Heaviside function and *κ* is the effective spring constant of the bond-linkers. When the arm is extending, the friction acts as a constant drag Γ_*i*_ = 1, while when the arm is retracting, the friction is due to the adhesion of the slip-bonds.

Eq.1 is a simplified description of the protrusive traction forces, which can be further elaborated to include the adhesion dependence of these forces [28]. The restoring force of the cell elasticity in Eqs.1,2 is described by a simple spring term (Fig. 1F(i)), where *k* is the effective elasticity of the cell (of rest length 1).

In Eq.2, *f*_*s*_ describes the susceptibility of the slip-bonds to detach due to the applied force, and *r* is the effective cell-substrate adhesiveness.

In Eq.3, *δ* is the rate at which the local actin flows relax to the steady-state solutions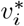, which is given by [27]

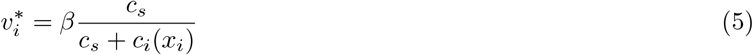

where *c*_*i*_(*x*_*i*_) is the concentration of actin polymerization inhibitor at the tip of arm *i* (at the coordinate *x*_*i*_), *c*_*s*_ is the saturation concentration, and *β* is the maximal actin polymerization speed at the arm edges. The term *σ ξ*_*t*_ in Eq.3 describes the noise in the actin polymerization activity, with a random Gaussian form of amplitude *σ*.

The calculation of the spatial distribution of the inhibitor concentration along the different branches *c*_*i*_(*x*_*i*_) was performed (Fig. S-1B(ii)). The spatial distribution is composed of exponential sections, maintaining continuity at the junctions, a no-flux boundary condition, and a constant total amount of inhibitor within the cell.

The total length of the cell is given by

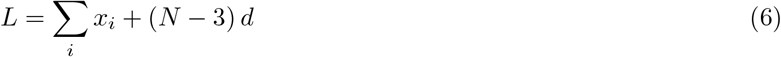

where *d* is the distance between two adjacent junctions on the network. Following the initialization of *x*_*i*_, *n*_*i*_ and *v*_*i*_ of each arm, their temporal evolution during the migration process can be obtained through numerical integration of Eq.1, Eq.2 and Eq.3. In this study, we employed the symmetric initial condition.

### Calculated shape dynamics of migrating cells

We analysed the shape dynamics of our model cell moving on a hexagonal network, as function of the actin polymerization speed parameter *β* and the grid size *d* (Fig. 2). We found that that migrating cells have more arms when *β* is larger, because the protrusive forces that elongate the cell are stronger, and longer cells span more junctions (Fig. 2A,B). Similarly, cells span more junctions and have more arms when *d* is smaller (Fig. 2A,B).

**Fig. 2:**
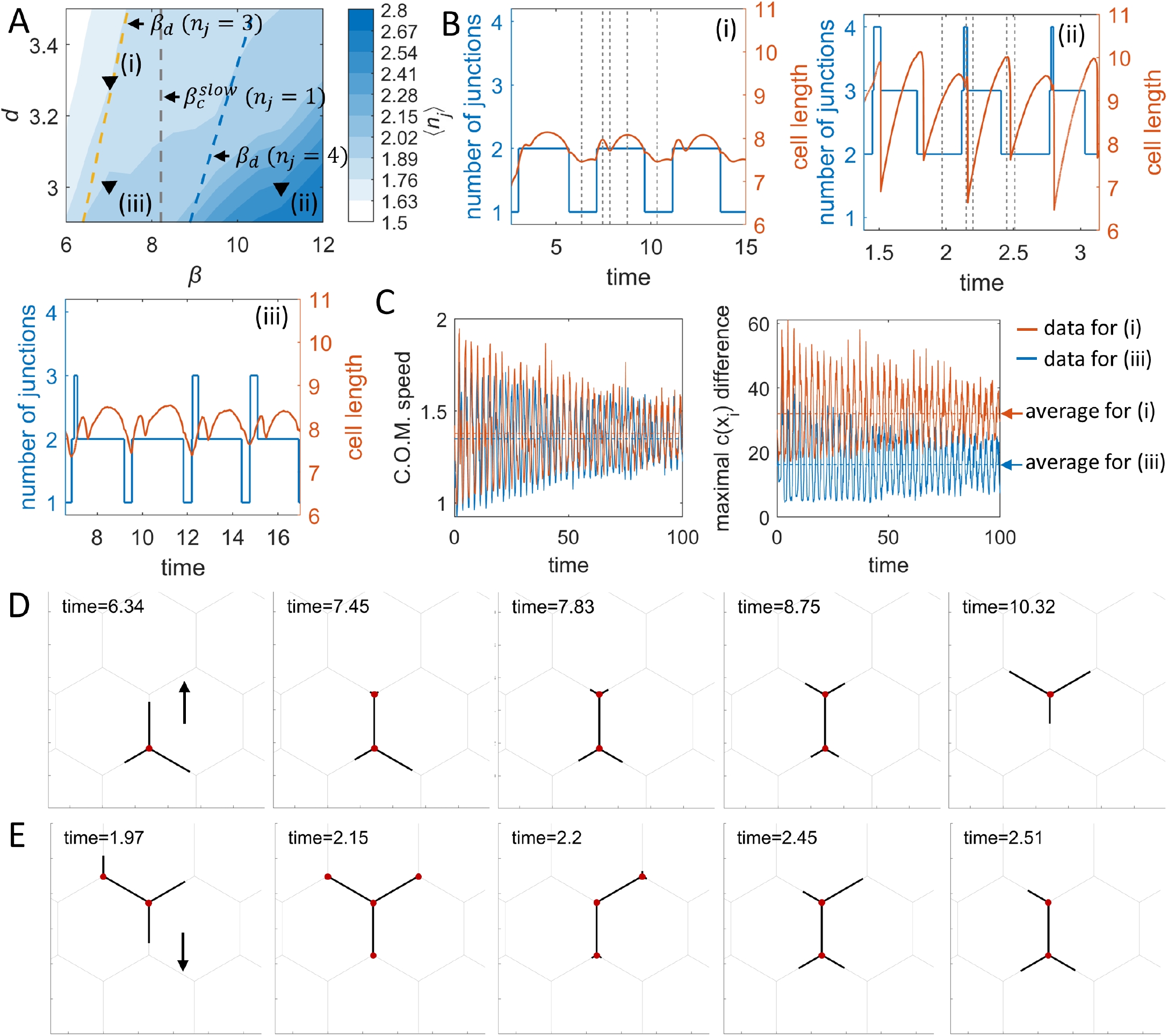
Simulations of facing multiple junctions during migration. (A) The *β−d* phase diagram of the time average of the number of junctions. Gray dashed line denotes the critical *β* of the slow process in the one-junction case. Yellow and blue dashed lines denote the critical *β*’s for cells to occupy 3 and 4 junction respectively, which are also the critical *β*’s for migration when cells are occupying 3 and 4 junctions (see Fig. S-1E, Eq. S-28). (B) The dynamics of the number of junctions and total cell length for sections (i-iii) in (A). (i) (*β, d*) = (7.0, 3.3). (ii) (*β, d*) = (11.0, 3.0). (iii) (*β, d*) = (7.0, 3.0). (C) Effect of number of arms on cell migration, comparing (i) and (iii) in (A): Left: Speed of the cells’ center-of-mass. Right: Maximal difference between the polarity cue concentration at the arm tips (*c*(*x*_*i*_)). (D) Snapshots for the time stamps (gray dashed lines) in B(i). (E) Snapshots for the time stamps (gray dashed lines) in B(ii). The arrows show the direction of migration. Parameters:*c* = 3.85, *D* = 3.85, *k* = 0.8, *f*_*s*_ = 5, *r* = 5, *κ* = 20, *δ* = 250, *σ* = 0.5.

We selected three regions from the phase diagram (black inverted triangles in Fig. 2A), to display the typical dynamics of the number of arms and the total cell length (Fig. 2B). By increasing *β* we observed that the number of junctions spanned by the cell is increased, due to the larger mean values and the larger fluctuations in the total cell length (Fig. 2B,D). Moreover, we found that the overall dynamics on the multiple junctions seems to be continuous as function of *β*, and they do not exhibit any sharp change in the behavior (Fig. 2A). This seems to be unique for multiple junctions, since for cells spanning a single junction there is a critical value of *β* above which the cell tends to form two equally persistent and long protrusions, that greatly increase the time the cell spends on the junction (the “slow process”) [17].

The critical *β* _*d*_, above which the cell extends to a length that spans *n*_*j*_ junctions, naturally increases with *n*_*j*_ (Figs. 2A, S-1E). We find that this critical *β* _*d*_ can be larger than the minimal value needed to allow the cell to polarize and leave the junction, *β* _*c*_ (See SI section S-2). When *β* _*c*_ *> β* _*d*_, as we demonstrate for cells spanning 2 junctions (Fig. S-1E), the cell can be trapped over the two junctions. The size of the cell that can be trapped while spanning *n*_*j*_-junctions depends on the model parameters (SI section S-2). This is due to the increased competition between the leading edges, which makes it more difficult for the cell to polarize as the number of the leading edges (arms) increase. The relation between the number of arms and migration is shown in Fig. 2C. We see that at the same value of *β*, and therefore very similar instantaneous cell speed and length (Fig. 2B(i),(iii), Movie S3), the cell polarity is significantly larger when the number of arms is smaller, on the hexagonal network of larger dimensions.

### Theoretical large-scale migration characteristics of cells

Next, we investigated the large scale migration characteristics of the cells on the hexagonal network. We calculated the mean-squared-displacement (MSD) of the center-of-mass (see SI section S-3) of the cells during migration. We plot the time series of *MSD* for different values of *β* and fit them by the power-law function *MSD* = *Kt*^*α*^ (Fig. 3A). We found that *Α ≈* 1 for all the sets, corresponding to regular diffusive motion. For this two-dimensional Brownian motion, the *MSD* is

**Fig. 3:**
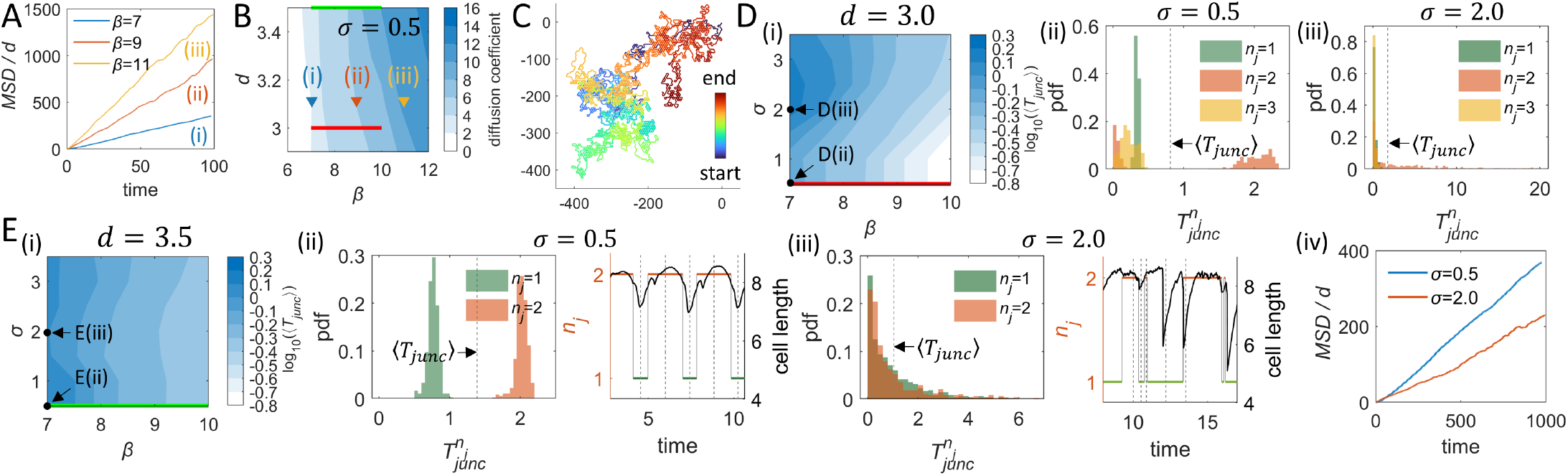
Analysis of the *MSD* of the cell’s centroid and junction residency time. (A) Time series of *MSD* for *β* = 7, 9, 11 and *d* = 3.1. (B) The *β − d* phase diagram of the diffusion coefficient of the cell’s centroid (*D*_*c*_, Eq.7). Red/Green line: the equivalent ranges of (*β, d*) values spanned by Red/Green line in D(i)/E(i). Maximal simulation time: *T* = 100. (C) A typical trajectory of cell’s centroid during a long run for *β* = 9 and *d* = 3.1. Maximal simulation time: *T* = 10000. Other parameters: *c* = 3.85, *D* = 3.85, *k* = 0.8, *f*_*s*_ = 5, *r* = 5, *κ* = 20, *δ* = 250, *σ* = 0.5. (D) (i) The *β − σ* phase diagram of *⟨T*_*junc*_*⟩* (in log scale) for *d* = 3.0. (ii) Distribution (probability density function, pdf) of 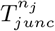for *β* = 7, *d* = 3.0, *σ* = 0.5. (iii) Distribution of 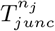 for *β* = 7, *d* = 3.0, *σ* = 2.0. (E) (i) The *β − σ* phase diagram of *⟨T*_*junc*_*⟩* (in log scale) for *d* = 3.5. (ii) Left: Distribution of 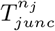 for *β* = 7, *d* = 3.5, *σ* = 0.5. Right: Time series of the number of junctions and total cell length for *β* = 7, *d* = 3.5, *σ* = 0.5. Snapshots for the time stamps (gray dashed lines) are shown in Fig. S-2C(i). (iii) Left: Distribution of 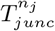 for *β* = 7, *d* = 3.5, *σ* = 2.0. Right: Time series of the number of junctions and total cell length for *β* = 7, *d* = 3.5, *σ* = 2.0. Snapshots for the time stamps (gray dashed lines) are shown in Fig. S-2C(ii). (iv) Time series of *MSD* for two different levels of noise, with *β* = 7 and *d* = 3.5.

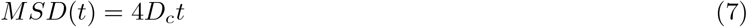

where *D*_*c*_ is the diffusion coefficient of the cell calculated by using the linear fit to the time series of the *MSD*. We find that *D*_*c*_ increases proportionally with *β* and *d* (Fig. 3B,C). This is expected, since in single junctions the residency time (which is defined as the time it takes a cell to leave a junction since it was first occupied) of cells at the junction generally decreases with increasing actin polymerization speed *β* [17]. Similarly, we study in Fig. 3D,E the junction residency time of the cell, *T*_*junc*_, as function of *β* and *σ* (which is the cellular internal noise, Eq.3). We used different *d* (*d* = 3.0 and *d* = 3.5) and found that the average residency time ⟨*T*_*junc*_⟩ decreases as *β* increases (Fig. 3D,E). This inverse relation is consistent with single junction case [17], in which the polymerization activity at the edges of the arms increases the migration speed, and decreases the residency time on a junction. This explains the role of *β* in increasing the cellular diffusion (Fig. 3B).

The effect of increasing the network dimension *d* is to increase the diffusion coefficient, due to decreasing the number of junctions that the cell is spanning (Fig.2A). By decreasing the number of junctions that the cell is spanning the polarity of the cell is stronger (Fig.2C), as there are fewer competing leading edges. The more robust polarity gives rise to more persistent migration, leading to higher diffusion coefficient.

For the network with *d* = 3.0 we found that the ⟨*T*_*junc*_⟩ increased as *σ* increases (Fig. 3D(i)). The noise increases the residency time as it decreases the strength of the flows that polarize the cell. We analysed the average residency time of the cell over a specific number of junctions, 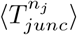,where *n*_*j*_ represents the number of junctions that the cell is spanning (Fig. S-2A). Cells spanning 2 junctions are especially stable at low noise (Fig. 3D(ii)). Two peaks with low and high 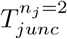 were observed as a consequence of the short time interval when *n*_*j*_ transits from 2 to 3, and the long time interval when *n*_*j*_ transits from 2 to 1 (Fig. 3D(ii)). At high noise, the residency time distribution becomes exponential, with a tail that reaches large values (Fig. 3D(iii)). This role of the noise as increasing the residency time was also found for cells on a single junction [17].

For a hexagonal grid of larger dimensions, *d* = 3.5, we find that moderate noise can decrease the average junction residency time (Fig. 3E(i)). This trend is driven by the dynamics of the 2-junction spanning cells (SI section S-4). Due to the larger grid size, the cells form shorter arms that extend at the ends of the spanned junctions, and therefore noise can drive these short arms to loose their “grip” on one of the junctions (Fig. 3E(ii,iii)). The corresponding simulation snapshots are shown in Fig. S-2C. This behavior is unique to cells spanning multiple junctions, whereby an increase in cellular noise decreases the time that a cell remain on a junction, and increases the frequency of hopping between junctions. However, the reduction of the junction residency time does not correspond to faster global cellular diffusion, as the *MSD* for high noise is significantly smaller than for low noise (Fig. 3E(iv)). The shorter junction residency time is driven by short bouts of the cell spanning two junctions and then loosing its “grip” on one of these junctions, a form of rapid oscillations around the same average location, and therefore do not correspond to faster overall migration. At higher values of noise we expect that the average junction residency time will eventually increase due to the loss of cellular polarization (Fig. 3E(i)).

An alternative way to quantify the migration over the network is the “hexagonal residency time” of the cells, *T*_*hex*_, which is defined as the time from when the cell’s center-of-mass enters a hexagonal cell of the network to when it leaves it and enter a new one. This quantification of the cellular migration is shown in the SI section S-5 and Fig. S-3.

### Comparison of the model with experiments

To compare our model with experiments, we performed live-cell imaging of HUVEC and bone marrow-derived macrophages cells on hexagonal networks of adhesive stripes (Fig. S-4), as the migration of both cell types is highly dependent on adhesion [23, 26]. We compared the calculated dynamics of the cell shape (number of junctions, cell length) as well as the distributions of junction residency times with the experimental observations. To extract these quantities from the experiments we adapted a pre-existing convolutional neural network image analysis [29, 30], to compute the positions of the tips of cellular arms that emanate from the junctions spanned by the cells.

Migrating HUVEC oscillated between spanning 1 and 2 junctions (Fig. 4A). Comparison with the simulations shows that the cells exhibit very similar dynamics of the total cell length, individual arm lengths, and the number of spanned junctions (Fig. 4B). Some HUVEC also exhibited frequent fluctuations between 1 and 2 junctions, or stable dynamics spanning 2 junctions (Fig. S-5A-D). To capture the variations observed between cells, we changed one of the cellular parameters in the model (since the variation in the micropattern dimensions are negligible, *∼* 1*™* 5%). We chose to vary the cell elasticity parameter *k*, since it affects the total length of the cell, and consequently the cell shape and the number of junctions it spans on the hexagonal network (Fig. 4B, Fig. S-5B,D). A similar change in dynamics arises if the hexagonal network has a slightly different size *d* (see simulation results in SI section S-4, Fig. S-2). The junction residency time distributions in both HUVEC experiments and simulations have a broad, exponential form (Fig. 4C,D). The comparison of the distributions from the simulations at different noise levels (Fig. 3D(ii,iii),E(ii,iii)) suggests that these cells are in a regime of large internal noise. Altogether, these data (Fig. 4A-D) shows good qualitative agreement between the model and the experiments in HUVEC.

**Fig. 4:**
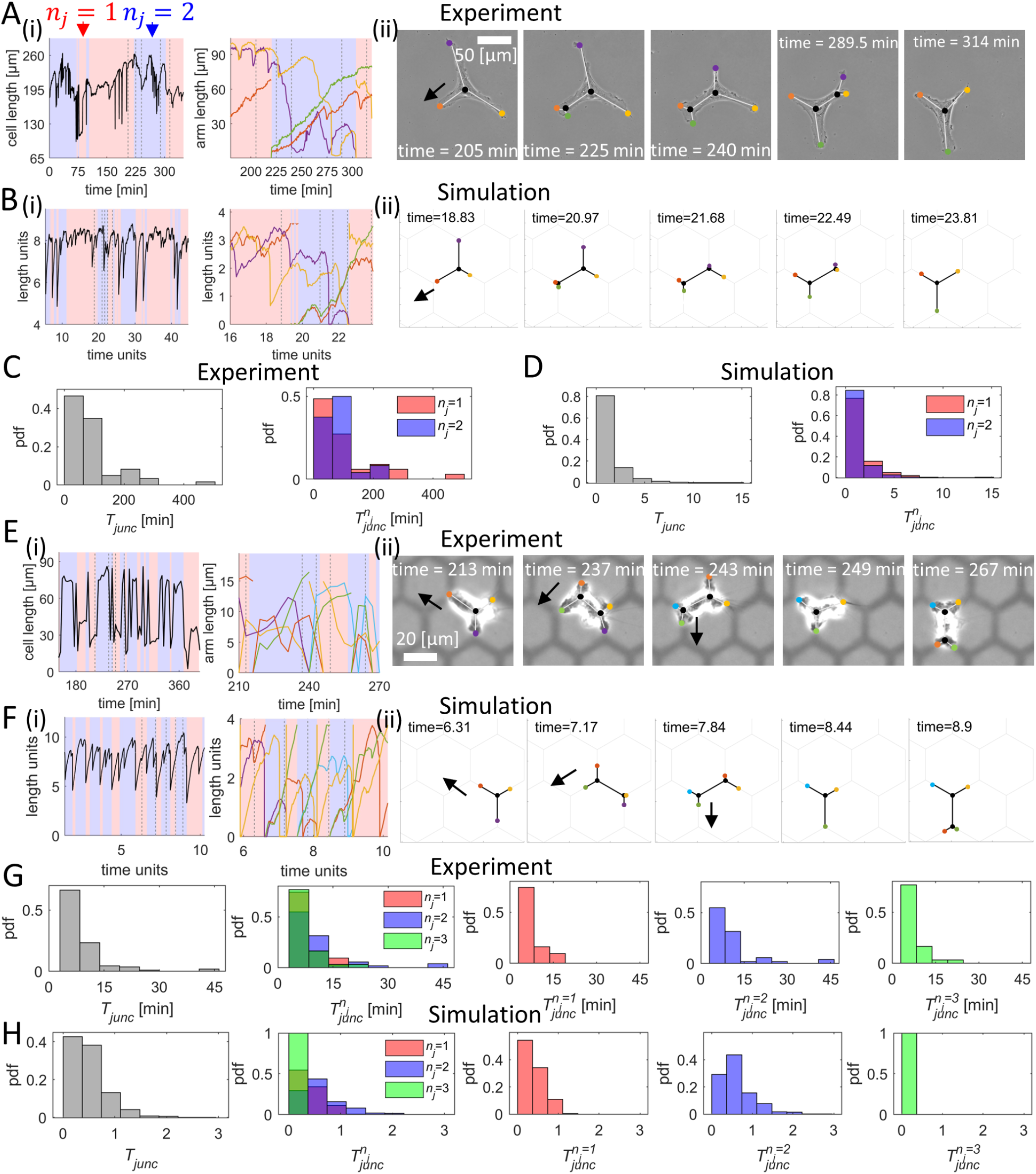
Comparisons between cell shapes and migration patterns in simulations and experiments on HUVEC and macrophages. (A) Experiments on migrating HUVEC (Movie S4). (B) Simulation of a migrating HUVEC (Movie S5). (i) Dynamics of the total cell length and the arms’ lengths. (ii) Snapshots of the cell, corresponding to gray dashed lines in (i). Key simulation parameters: (*β, d, σ*) = (7.0, 3.6, 2.0). Other simulation parameters: *c* = 3.85, *D* = 3.85, *k* = 0.8, *f*_*s*_ = 5, *r* = 5, *κ* = 20, *δ* = 250. (C) Distribution (pdf) of *T*_*junc*_ (left) and distribution of 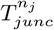 (right) in the HUVEC experiments. 10 cells across 5868.5 minutes were used to obtain these distributions. (D) Distribution of *T* (left) and distribution of 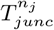 (right) from a long simulation with the parameters of (B). (E) Experiments on migrating macrophages (Movie S6). (F) Simulations of migrating macrophages (Movie S7). i) Dynamics of the total cell length and the arm lengths. ii) Snapshots of the cell, corresponding to gray dashed lines in (i). Key simulation parameters: (*β, d, σ*) = (10.0, 3.8, 3.0). Other simulation parameters: *c* = 3.85, *D* = 3.85, *k* = 0.8, *f*_*s*_ = 5, *r* = 5, *κ* = 20, *δ* = 250. (G) Distribution (pdf) of *T*_*junc*_ (leftmost) and distribution of 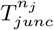 (right ones) in the macrophages experiments, where the cells spans 1, 2 or 3 junctions. A total of 2 cells across 849 minutes were used to obtain these distributions. (H) Distribution of *T*_*junc*_ (leftmost) and distribution of 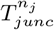 (right ones) from a long simulation with parameters of (F).

Macrophages efficiently explored the hexagonal lattice, as the number of junctions spanned oscillated between 1 and 3. In comparison to HUVEC, macrophages exhibited a shorter junction residency time, with a narrow distribution (compare Figs. 4C,D and 4G,H). This suggests that these cells are in a regime of higher overall actin protrusive activity (the parameter *β* in the model) and lower internal noise (Fig. S-5E,F). These properties lead to a higher polarization of macrophages in comparison to HUVEC, which is expected for immune cells that need to efficiently migrate in complex microenvironments [20, 31].

We next compared the results of simulations and three experiments of HUVEC that are stuck at the junctions (Fig. S-6), and are not globally motile over the timescale of the experiment. In our simulations this behavior occurs when the internal noise is too large for cells to efficiently polarize and migrate away from the junction. In the first example (Fig. S-6A,B), the number of junctions oscillates over the range 1 to 3. In the second and third examples (Fig. S-6C-F), the number of junctions oscillates between 2 and 3.

Overall these observations are consistent with our model: cells, such as HUVEC, that exhibit shape dynamics that correspond to lower actin activity (*β*) and higher internal noise, also exhibit lower migration and higher tendency to get trapped on junctions, especially when spanning more than one junction. Cells that correspond to higher actin activity, such as macrophages, exhibit efficient migration even when having highly bifurcated shapes.

We next tested the effect of substrate adhesion in our model. Mesenchymal migrating cells require firm adhesion to the substrate by forming focal adhesions, which are then disassembled at the cell rear by calpain (a *Ca*^2+^-dependent protease) [1, 20, 25]. To increase cell adhesion we used an inhibitor of calpain (50 µM PD150606). As expected calpain inhibition increased the substrate adhesiveness, and the treated HUVEC had elongated shapes (Fig. 5A). Moreover, inhibition of focal adhesion disassembly slowed the cell rear detachment and the elongated cells spanned a larger number of junctions, oscillating between 2 and 4 (Fig. 5A). Similar results were obtained in the simulations by increasing the adhesion strength (*r*) and both the overall actin polymerization activity (*β*) and its noise (*σ*), compared to the normal HUVEC cells in Fig.4. These changes have the effect of increasing the cell length and number of spanned junctions (Fig. 5B). Consequently, experiments in HUVEC and simulations show similar patterns of the junction residency time, with relatively short residency times for both *n*_*j*_ = 2, 4, and the widest distribution extending to long residency times for *n*_*j*_ = 3.

**Fig. 5:**
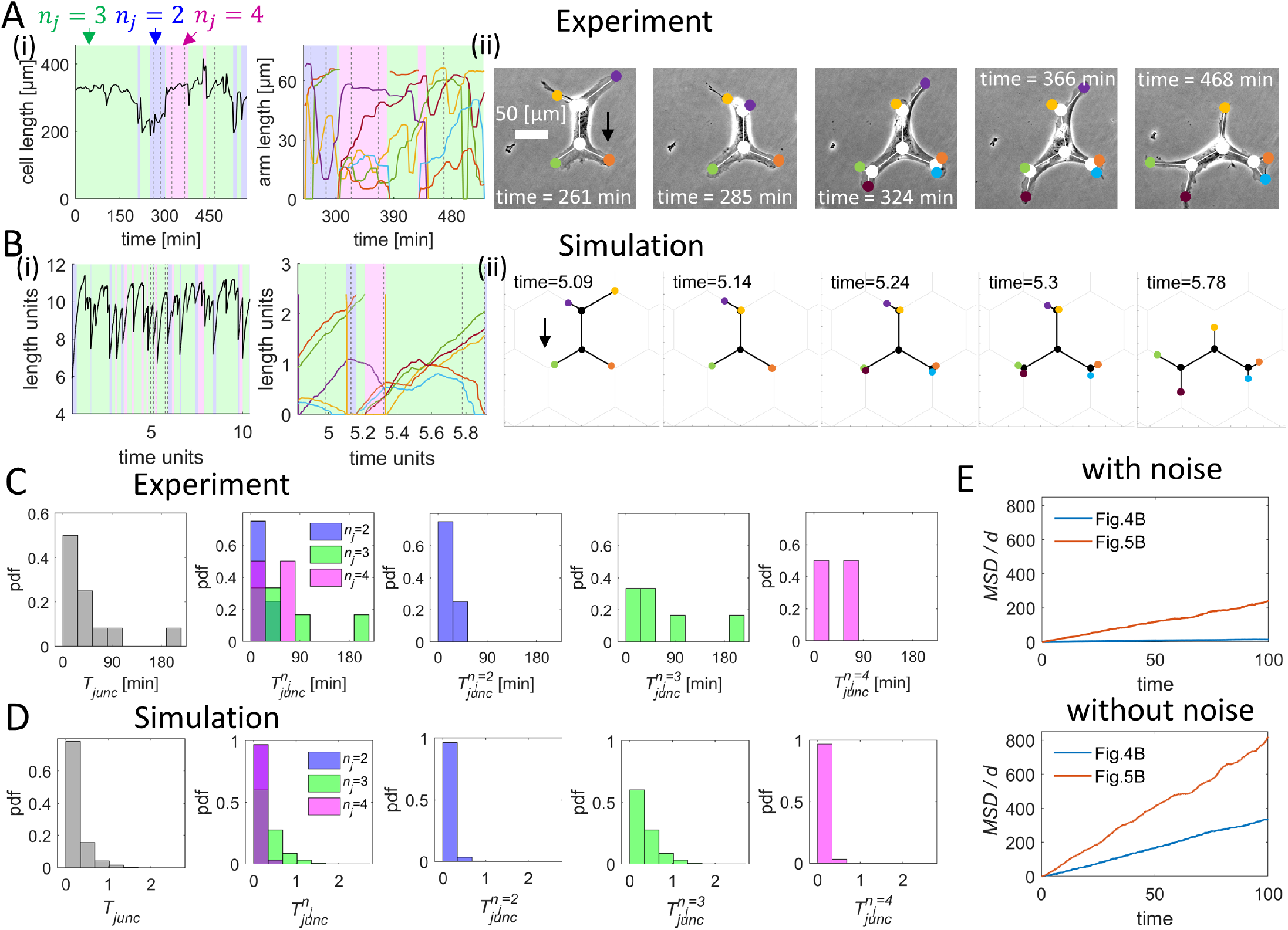
Comparisons between cell shapes dynamics in simulations and experiments for HUVEC with larger cell-substrate adhesiveness. (A) Experiments on HUVEC treated with 50 µM PD150606, a calpain inhibitor (Movie S8). (B) Simulations of HUVEC with larger cell-substrate adhesiveness (Movie S9). Hexagon side: 60 µm. Inter-hexagon width: 20 µm. i) Dynamics of the total cell length and the arm lengths. ii) Snapshots of the cell, corresponding to gray dashed lines in (i). Key simulation parameters: (*β, d, σ, r*) = (10.0, 2.4, 2.7, 7.0) in (D). Other simulation parameters: *c* = 3.85, *D* = 3.85, *k* = 0.8, *f*_*s*_ = 5, *κ* = 20, *δ* = 250. (C) Distribution (pdf) of *T*_*junc*_ (leftmost) and distribution of 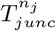 (right ones) in the experimental regime in which the cells spans 2, 3 or 4 junctions. One cell across 576 minutes was used to obtain these distributions. (D) Distribution of *T*_*junc*_ (leftmost) and distribution of 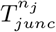 (right ones) from a long simulation with parameters of (B). (E) Comparisons of *MSD* of the simulated HUVEC of Fig. 4 and Fig. 5, with the noise levels identical to those used in the corresponding simulations (top) or without the noise (bottom).

In Fig. 5E we show the calculated effects of the changes induced by the drug on the large-scale diffusion of the simulated cells. We compared the normalized *MSD* of the HUVEC cells of Figs. 4B,5B, with and without the actin polymerization noise (see also Fig. S-7C). As expected the diffusion decreases for lower *β* and higher adhesion, even without noise, due to the cell speed decreasing. However, large noise has an additional dominant effect that can lead to long residency times, due to getting trapped over multiple junctions, and therefore result in a drastic decrease of the diffusion coefficient *D*_*c*_ (Eq.7) over the network.

Finally, and to provide an *in vivo* validation of our model we analyzed the migration of interstitial macrophages moving in the zebrafish tail fin. We find that the cells have irregular branched shapes (Fig. 6A), as they branch along the regular array formed by basal keratinocytes that have a polygonal shape [32]. Due to the cellular heterogeneity expected in a tissue, the shapes have different lengths of the sections between junctions (*d*, Fig. 6B), unlike the regular hexagonal networks of the *in-vitro* experiments and simulations. However, the distances between the junctions are within a similar range of values compared to that used in the *in vitro* experiments. Similar to the observations made in the hexagonal micropatterns, the junction residency time shows a tight peak (Fig. 6C). The relation between the number of junctions and the cell length is in good agreement with the data obtained in both experiments and simulations (Fig. 6D), showing that our model can be used to describe the shape dynamics and migratory behaviour of macrophages *in vivo*.

**Fig. 6:**
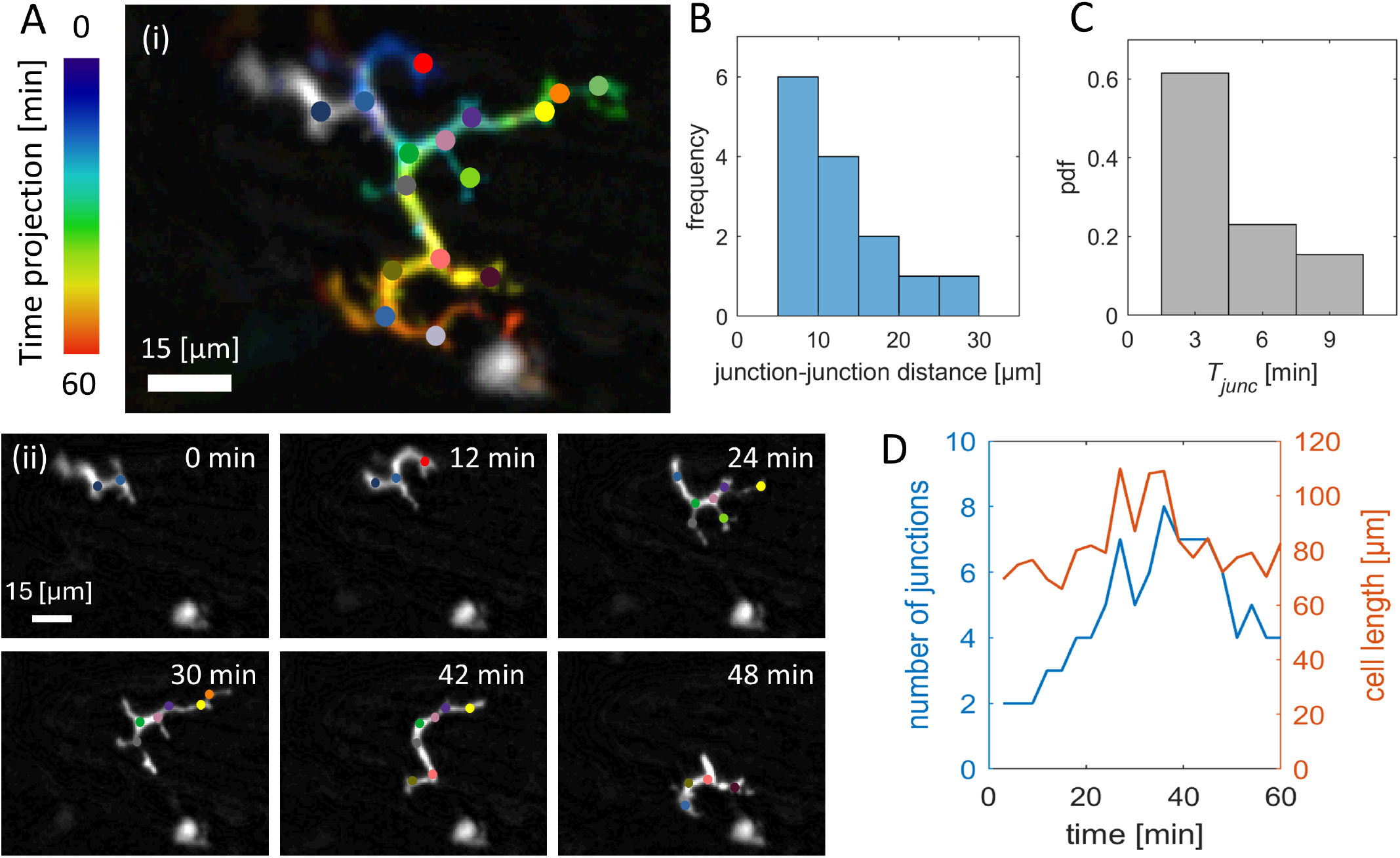
Analysis of *in vivo* migration. (A) (i) Temporal color code showing the trajectory of a macrophage migrating in the zebrafish tailfin. Junctions are depicted with colored dots. This label is the same for panels (i) and (ii). (ii) Snapshots of a migrating macrophage at indicated times. (B) Distribution of the distance between each pair of adjacent junctions. Note that *in vivo* this parameter is variable. (C) Distribution of the macrophage’s residency time (*T*_*junc*_). (D) Dynamics of the number of junctions and total cell length across time.

## Discussion

In this work, we presented a theoretical model, and the corresponding *in vitro* and *in vivo* experimental proof of concept, for the shape dynamics and migration of branched cells inside complex networks. The model describes the self-polarization of the cell due to the feedback between the actin polymerization activity at the leading edges of the cellular protrusions, and the global redistribution of a polarity cue that acts to inhibit local actin polymerization and protrusive activity. Here, we describe migrating cells that span multiple junctions and have a highly ramified shape with multiple competing arms in geometries such those observed during cellular migration within tissues *in vivo*[20, 31]. This model extends our recent description of cellular directional decision-making over a single junction [17], and it reveals an unknown trade-off between microenvironmental exploration for directional cues and long-range migration efficiency during directional decision-making.

We compare the cellular dynamics predicted by the theoretical model with the migratory behaviour of two cell types that undergo stick-slip migration: HUVEC and macrophages, which use a mesenchymal migratory mode [1, 23, 25, 26]. These cells were allowed to migrate on a hexagonal network of adhesive lines, which resemble tissue geometries. The network dimensions were adapted to each cell: by constraining the cells to simultaneously span several junctions we trigger a high level of branching. The model provides a good qualitative description of the observed cellular shape dynamics.

Using our model we investigated the role of noise in the polymerization activity, and the results suggest that cells with overall weak and very noisy internal actin retrograde flows, are susceptible to losing their polarity and getting trapped at the network junctions. This indicates that there is an optimal range for the number of protrusions during cell migration: On the one hand these protrusions contribute to pathfinding and direct cellular migration by allowing the cells to better sense the microenvironment. On the other hand, if the number of protrusions is too high this will hinder efficient migration. This trade-off between microenvironmental sensing and efficient motility may determine the optimal levels of acto-myosin contractility and protrusive activity, for each cell type. Our data suggests that this trade-off evident during the faster migration of macrophages in the hexagonal array, in comparison to HUVEC. This could be directly related to the surveillance function of macrophages, which requires the efficient exploration of the microenvironment, while preserving their migratory capacity [31].

Acto-myosin contractility requires *Ca*^2+^ signals in the cytoplasm. Interestingly, *Ca*^2+^ signaling proteins are directly involved in migration and branching ([7]). The latter suggest a prominent role for calcium signaling during this response. Consequently, the manipulation of calcium-dependent protease calpain lead to an increase in the cell length and to a less efficient directional decision-making. In addition, we found that the variations in the total cell length were higher in the more motile macrophages than in HUVEC. These findings might have implications regarding cell volume control, which also impacts cell migration [33–36]. Fast migrating macrophages would then require equally faster adaptation to the microenvironment, and therefore need rapid changes in the function of molecules involved in cell volume control. Therefore, in macrophages we anticipate a prominent role for different cell volume regulators, such as mechanosensitive channels, tranporters/exchangers, and pumps [33–35], which might contribute to provide the shape plasticity necessary for the efficient migration of these cells. Ion channels may also be essential for a mechanism that cells can employ to release themselves from being trapped at junctions, and to restart motility, which is to undergo a calcium-mediated “reset” of the actin cytoskeleton [37]. Future studies should be performed to evaluate these hypotheses.

The model presented in this work provides us with the framework to explore in the future the complex migration of bifurcated cells at the subcellular level, to reveal the details of the cytoskeletal activity during this process. This model extends the current conceptual and predictive frameworks [17, 19, 27] to study cell migration inside 2D and 3D networks composed of linear tracks linked in different complex geometries. In addition, the present model can allow to study how such geometries affect the ability of cells to perform chemotaxis and haptotaxis in response to external gradients [38]. This will be most relevant for cells migrating through dense tissues, while guided by external signals, such as immune [21], cancer [2] cells, and others.

### Experimental Methods Cell culture

#### Macrophages

Bone marrow derived macrophages (BMDMs) were differentiated from myeloid precursors to macrophages in the span of 7 days. Briefly, bones were collected under protocol ORG 1062 granted to Prof. Pablo J. Sáez by the Animal Welfare Officers of the University Medical Center Hamburg-Eppendorf (UKE). Bone-marrows were isolate as previously described [39], and the precursors were cultured as it follows. Briefly, precursors were isolated and plated in Corning culture dishes of 100 [mm] diameter using RPMI 1640 Medium, GlutaMAX™ Supplement, HEPES (10% FCS) (Gibco, ThermoFisher), supplemented with 1% Penicillin-Streptomycin (PenStrep, ThermoFisher) and 100 [µg/mL] of Macrophage Colony-Stimulating Factor (Bio-Techne GmbH). The cells were incubated at 37°C and 5% CO2 for 7 [days], adding 5 [mL] of RPMI 1640 Medium, GlutaMAX™ Supplement, HEPES (10% FCS), supplemented with 1% Penicillin-Streptomycin, and 100 [µg/mL] of Macrophage Colony-Stimulating Factor (M-CSF) on days 3 and 5. On day 7, cells were detached using ethylenediaminetetraacetic acid (EDTA) 5 [mM] (ThermoFisher), and subsequently plated in the micropatterns.

#### HUVEC

The culture of Human Umbilical Vein Endothelial Cells (HUVEC) was performed as described previously [17]. Briefly, HUVEC were acquired from PromoCell (C-12203) were used between passages 5 and 12. These cells were originally isolated from the vein of the umbilical cord of pooled donors. The HUVEC were cultured in full endothelial growth medium that included the Basal Medium and the SupplementMix (PromoCell), supplemented with 1% Penicillin-Streptomycin (PenStrep, ThermoFisher), on 100 [mm] diameter Corning culture dishes, coated with fibronectin from bovine plasma (1 [µL/mL]) for 20 [min] at 37°C and 5% CO2. The dishes were washed once with Dulbecco’s Phosphate-Buffered Saline (DPBS, 1X, Gibco, ThermoFisher) prior to the culture. Once cultured, the cells were incubated at 37°C and 5% CO2 for 2 [days] until reaching a confluency of 80%. Once the monolayer is formed, the cells were passaged by detaching the cells with 3 [mL] of TrypLE Express (with Phenol Red, 1X, Gibco, ThermoFisher) for 2 [min] and then plating 1,5×106cells/[mL] in a new coated Corning culture dishes with 100 [mm] diameter in a total of 10 [mL] new full endothelial growth medium.

#### Micropatterning

The micropatterning was performed using the photopatterning technique, possible with a Digital Micromirror Device (Primo™, ALVEOLE) coupled to a T2i Eclipse microscope (Nikon). A Polydimethylsiloxan (PDMS) stencil with a circle area of 5 [mm] was placed inside a bottom glass dish (FluoroDish, World Precision Instruments), which is a 35 [mm] diameter glass bottom dish, previously cleaned with a PlasmaCleaner (PDC-32G-2, Harrick Plasma) for 5 [min]. Afterwards, the area within the stencil was coated for 1 [hr] at room temperature with 1 [mg/mL] pLL-PEG, in the case of the HUVEC, or fluorescent PEG atto 633 (SuSoS AG), in the case of macrophages. For HUVEC and Macrophages, the remaining pLL-PEG was washed with sterile water. Then in both cases, 10 µL of photo-activator (PLPP™, ALVEOLE) was added, and then degraded using UV illumination at a power of 600 [mJ/mm2], with the shape of the desired pattern. The pattern consisted of a hexagonal array, with a side of 60 [µm] and a width between hexagons of 20 [µm] in the case of HUVEC; and a side of 12 [µm] and a width between hexagons of 4 [µm] in the case of macrophages. These patterns were created using the software Inkscape 1.0.2-2 (see SI section S-6, Fig. S-4). For HUVEC, the photo-activator (PLPP™, ALVEOLE) was washed with 1X DPBS and then the dish was coated for 20 min at 37°C with fibronectin (1 [µL/mL]) mixed with fibrinogen (1:10) to visualize the patterns, and then rinsed with 1X DPBS. For macrophages, the remaining photo-activator (PLPP™, ALVEOLE) was washed out with 1X DPBS and where left uncoated to let the cells adhere to the glass of the bottom of the dish. After this 1X DPBS was removed and replaced with the corresponding media. Finally, the cells were plated as it follows. 10.000 HUVEC were seeded and left overnight (around [16h]) at 37°C and 5% CO2 to ensure their adhesion to the plate. In the case of macrophages, 10,000 cells were plated and imaged around 6-8 [hr] later.

#### Transfection

HUVEC were cultured until reaching a confluence of 80%, and detached as mentioned above in the section for cell culture. Then the cells were transfected using the 4D-NucleofectorTM X Unit (Lonza), with 1 [µg] of F-tractin GFP plasmid (Plasmid #58473 Addgene) and the P5 Primary Cell 4D-NucleofectorTM X Kit to then electroporate corresponding to the transfection program (CA-167) to then keep overnight (around 16 [hr]) in culture previous to use.

#### Live-cell imaging

The imaging for the phase contrast movies was performed using an inverted microscope (Leica Dmi8) equipped with an APO 10x/0.45 PH1, FL L 20x/0.40 CORR PH1, and APO 40x/0.95 objectives. Images were recorded with an ORCA-Flash4.0 Digital camera (Hamamatsu Photonics) using the MetaMorph Version 7.10.3.279 software (Molecular Device). The actin fluorescent videos were performed with a Nikon Eclipse TiE equipped with a 40x Plan Fluor Phase objective. Images were recorded with an Photometrics Prime 95B (back-illuminated sCMOS, 11 µm pixel-size, 1200×1200 pixels) connected to a Yokogawa CSU W-1 SoRa in dual-camera configuration c onnected to a Confocal disk with 50 µm pinholes using the VisiView v4 software. During every acquisition, cells were kept at 37°C and 5% CO2. For HUVEC, the acquisition was done with the FL L 20x/0.40 CORR PH1, actin visualization was done with 40x Plan Fluor Phase objective of the Nikon Eclipse TiE, both where imaged with a time interval of 3 [min], the rest of the phase contrast movies where imaged with the APO 40x/0.95 objective with a time interval of 30 [sec]. For macrophages, the acquisition for movies was done with an APO 40x/0.95 objective, with a time interval of 3 [min]. Treatment with 50 µM PD150606 calpain inhibitor was added just before acquisition.

#### Tracking and neuronal network

The coordinates of the tips of the cells across the junctions were monitored using neural networks with the ResNet50 architecture described previously [29, 30]. The were trained to analyze the movement of the cells and extract these coordinates. For labelling the movies, different frames were extracted from the movies in regular quantities of 12 units. This means that if the video was 120 frames long, the 120 would be divided by 12 and the frames would be extracted 10 by 10 accordingly. Labels were added manually in both the tips of the cells and the junctions, in each of the extracted frames. For HUVEC, a unique neural network was developed for all the movies as the similarities between the movies facilitated the generalisation of the analysis, training a network on the labelled data for 175,000 iterations. For macrophages, neural networks were developed per single movie, and each neural network was trained during 300,000 iterations on the labelled data, which provided a reliable basis for each macrophage movie analyzed. The data provided by the neural networks consist of three columns per label (X and Y position, and the likelihood of the prediction) and a row per frame. This data was represented in a video using a cutoff for the likelihood of 0.6, supervising the predictions and correcting them by hand when the prediction was not in the tip of the cell. Also, the predictions for the position of the junctions, as they were a fixed point, were substituted by the exact position X and Y of each junction. To obtain the junction residency time, the following steps were followed. Frame by frame, the number of junctions in which the cell was spanning was annotated. With this information, the frequency of the amount of frames in which the cell was spanning a certain number of junctions was extracted. Finally, these distributions of the junction residency time were compared with those obtained from the simulation data.

#### Zebrafish *in vivo* experiments

The zebrafish (*Danio rerio*) for the imaging experiments were kept in a UK Home Office-approved facility. The standards of the Animal (Scientific Procedures) Act UK 1986 were followed for the experiments. The zebrafish larvae were between 3 days post-fertilisation (dpf) and 5 dpf for the experiments.

The transgenic line Tg(mpx:EGFPi114; MPEG1:mCherry) was used to monitor the migratory behavior of macrophages [40]. Briefly, adult fish were kept at 28.5°C in a 14:10 hour light:dark cycle. Eggs were collected and kept in 90mm petri dish with methylthioninium chloride (methylene blue at 0.5 mg/mL). The fertilised eggs were also kept at 28.5°C until the end of the experiment.

Imaging was performed using a 6 well-plate in which the larvae were kept anaesthetised with 4.2% (v/v) tricaine and embedded in agarose. The 1.5% agarose was prepared and let to cool before mounting the larvae to avoid any damage to the larvae. Once the larvae were mounted and the agarose was set the well was filled with methylene blue and 4.2% (v/v) tricaine to avoid dehydration throughout. The plate was placed in the stage of the EVOS FL Auto2 and the humidifier function was used to set the temperature to 28.5 °C. The 10x magnification was used for the time-lapse function, where an image was taken every 3-5 minutes over the course of 20-24 h. The images were acquired every 3 min using brightfield, as well as GFP and Texas red filters in the microscope.

## Author contributions

Conceptualization: N.S.G. and P.J.S. Methodology: J.L., J.E.R. and N.S.G. Software: J.L. performed the model simulations and J.B.-C. designed the image analysis pipeline for the experimental data. Formal analysis: J.L. and J.E.R. Investigation: J.M.K. performed all the migration experiments in HUVEC, and with J.B.-C. performed the migration experiments in macrophages. M.E.M.O. performed all the analysis in zebrafish. Resources: A.G.R. and P.J.S. Writing - Original Draft: All authors. Writing - Review and Editing: All authors. Visualization: J.L. and J.B.-C. Supervision: A.G.R. supervised the *in vivo* experiments, N.S.G. supervised the model simulations and P.J.S. supervised the *in vitro* experiments. Project administration: N.S.G. and P.J.S. Funding acquisition: A.G.R., N.S.G. and P.J.S.

