## Supplementary materials for "Trade-off between branching and polarity controls decision-making during cell migration"


### S-1. Steady-state actin treadmilling flows and polarity cue distribution profile

#### Calculation of the actin treadmilling flows

We assume that the local actin treadmilling flow emanating from the free edge of an arm splits symmetrically at each junction. For example, for a cell spanning 2 junctions, the flow from arm 1, $v_1$, splits symmetrically into arm 2 and the segment between node P and node Q (Fig. S-1A(i)). Considering $v_i$ of all the arms, the net actin treadmilling flow $u_i$ within each arm is given by the sum of the flows that enter that arm

$$
\begin{aligned}
u_1 &= v_1 - \frac{v_2}{2} - \frac{v_3}{4} - \frac{v_4}{4} \\
u_2 &= v_2 - \frac{v_1}{2} - \frac{v_3}{4} - \frac{v_4}{4} \\
u_3 &= v_3 - \frac{v_4}{2} - \frac{v_1}{4} - \frac{v_2}{4} \\
u_4 &= v_4 - \frac{v_3}{2} - \frac{v_1}{4} - \frac{v_2}{4}
\end{aligned} \tag{S-1}
$$

with the convention that a positive flow is away from the arm tip (Fig. S-1A(ii)). The middle segment that connects the two adjacent junctions P and Q has a net flow from all the incoming segments that connect to it

$$u_m = \frac{v_3}{2} + \frac{v_4}{2} - \frac{v_1}{2} - \frac{v_2}{2} \tag{S-2}$$

and we take it to be positive towards junction P. Generally, the net actin flow of a free arm $i$ is given by

$$u_i = v_i - \sum_{j \neq i} \frac{v_j}{2^{m_{i,j}}} \tag{S-3}$$

where $m_{i,j}$ is the number of the junction nodes between arm $i$ and arm $j$. Note that the relation $\sum_{j \neq i} \frac{1}{2^{m_{i,j}}} = 1$ always holds.

The net actin flow of a node connection segment $k$ that connects two adjacent junction nodes Q and P is given by

$$u_k = \sum_{i\ connect\ to\ Q} \frac{v_i}{2} - \sum_{j\ connect\ to\ P} \frac{v_j}{2} \tag{S-4}$$

where $\{i\}$ and $\{j\}$ denote the free arms that emanate from junction Q and P, respectively. The positive flow of $u_k$ is taken to be along the direction from Q to P.

Note that within the cell, due to the nucleus and other organelles, the actin does not freely flow along the entire length of the cell, as suggested by our model (Fig. S-1A). However, these flows drive the formation of an inhomogeneous concentration profile of the cytoplasmic contents (proteins as well as larger organelles) along the cell length, and this is captured by our simplified model.

**Concentration profile of the polarity cue (actin polymerization inhibitor)**

We consider a polarity cue $c(x)$ diffusing in the cytoplasm, which is advected by the net actin treadmilling flows and acts as the inhibitor of actin polymerization. We assume that the concentration profile of the polarity cue relaxes faster than the timescale of the changes in the actin flow or cell shape [1–3]. This assumption greatly simplifies the problem and allows us to use the steady-state distribution for this concentration field, which is updated as the actin flows and cell shape changes. In the steady state, the advection-diffusion equation of the polarity cue within each arm is

$$\frac{\partial}{\partial x}(u_i\, c_i(x) + D\,\frac{\partial c_i(x)}{\partial x}) = 0 \tag{S-5}$$

where $u_i$ is the net actin flow in arm $i$, $c_i(x)$ is the concentration of the polarity cue and $D$ is the diffusion coefficient of the polarity cue. This gives a solution

$$c_i(x) = c_{i,0}\, \exp\left(-\frac{u_i x}{D}\right) + c_{i,1} \tag{S-6}$$

By applying a no-flux boundary condition at the ends of the arms, i.e., $u_i c_i(x) + D\frac{\partial c_i(x)}{\partial x}|_{x=x_i} = 0$, we obtain the coefficient $c_{i,1} = 0$. The relationship between the coefficients $c_{i,0}$ $(i = 1, 2, ..., N)$ is calculated based on the continuity of the concentrations at the junction nodes, and it depends on the cell shape. For example, for a cell spanning 2 junctions, we denote node P as the reference point of polarity cue concentration with the concentration as $c_0$ (Fig. S-1B(i)), and then the concentration profiles for all the arms and the middle segment can be written as

$$\begin{aligned} c_{1/2}(x_{1/2}) &= c_0\, \exp\left(-\frac{u_{1/2}\, x_{1/2}}{D}\right) \\ c_m(x_m) &= c_0\, \exp\left(-\frac{u_m\, x_m}{D}\right) \\ c_{3/4}(x_{3/4}) &= c_0\, \exp\left(\frac{-u_m\, d}{D}\right)\, \exp\left(-\frac{u_{3/4}\, x_{3/4}}{D}\right) \end{aligned} \tag{S-7}$$

The corresponding graphs are showed in Fig. S-1B(ii).

The total concentration of the polarity cue is conserved within the cell

$$c_{tot} = \int_0^d c_m(x_m)\; dx_m + \sum_{j=1}^{4} \int_0^{x_j} c_j(x_j)\; dx_j \tag{S-8}$$

where the sum over $\{j\}$ represents the contributions of the free arms. By substituting Eq.S-7 into Eq.S-8, we obtain $c_0$ as

$$\begin{aligned} c_0 = \frac{c_{tot}}{D} \Bigg[ & \frac{1 - \exp\left(-\frac{u_m d}{D}\right)}{u_m} + \frac{1 - \exp\left(-\frac{u_1 x_1}{D}\right)}{u_1} + \frac{1 - \exp\left(-\frac{u_2 x_2}{D}\right)}{u_2} \\ & + \exp\left(-\frac{u_m\, d}{D}\right) \left( \frac{1 - \exp\left(-\frac{u_3 x_3}{D}\right)}{u_3} + \frac{1 - \exp\left(-\frac{u_4 x_4}{D}\right)}{u_4} \right) \Bigg]^{-1} \end{aligned} \tag{S-9}$$

Generally, one can arbitrarily choose the reference point of the concentration field, denoted by $c_0$, at one of the junctions. Then the concentration in segment $i$ ($i$ could be either the free arm or the node connection segment) can be written as

$$c_i(x) = c_0\, \exp\left(-\frac{\sum_k u_k\, d}{D}\right)\, \exp\left(-\frac{u_i x}{D}\right) \tag{S-10}$$

where $\{k\}$ denotes the node connection segments between the segment $i$ and the reference point. The direction of $u_k$ is towards the reference point.

The total amount of the polarity cue is conserved within the cell

$$c_{tot} = \sum_l \int_0^d c_l(x_l)\, dx_l + \sum_j \int_0^{x_j} c_j(x_j)\, dx_j \tag{S-11}$$

where $\{l\}$ represent the nodes connected by full segments (of length $d$) and $\{j\}$ represent the free arms. From Eqs.S-10,S-11 the value of $c_0$ can be found

$$c_0 = \frac{c_{tot}}{D}[\sum_l exp(-\frac{\sum_{k,l} u_{k,l}\, d}{D})\frac{1-exp(-\frac{u_l\, d}{D})}{u_l} + \sum_j exp(-\frac{\sum_{k,j} u_{k,j}\, d}{D})\frac{1-exp(-\frac{u_j\, x_j}{D})}{u_j}]^{-1} \tag{S-12}$$

where the summation $k, l/j$ is for the nodes connected by segments $\{k\}$ that are between the segment $l/j$ and the reference point.

By substituting the expression of the inhibitor concentration (Eq.S-10) into the expression of the steady-state local actin flows (Eq.5), we obtain an implicit equation group of $v_i$ $(i = 1, 2, ..., N)$

$$v_i = \beta \frac{1}{1 + c_0\, exp(-\frac{\sum_k u_k\, d}{D})\, exp(-\frac{u_i x}{D})} \tag{S-13}$$

where $u_i$, $u_k$ and $c_0$ is given by Eq.S-3, Eq.S-4 and Eq.S-12, respectively. By numerically solving Eq.S-13, we can determine the values of $v_i$ corresponding to a specific set of $\{x_i\}$.

For the 2-junction case, the equation group of $v_i$ $(i = 1, 2, 3, 4)$ is obtained by substituting Eq.S-7 into Eq.5

$$\begin{aligned} v_{1/2} &= \beta(1 + \frac{c}{D}\frac{exp(-\frac{u_{1/2}x_{1/2}}{D})}{\frac{1-exp(-\frac{u_m\, d}{D})}{u_m} + \frac{1-exp(-\frac{u_1\, x_1}{D})}{u_1} + \frac{1-exp(-\frac{u_2\, x_2}{D})}{u_2} + exp(-\frac{u_m\, d}{D})\,(\frac{1-exp(-\frac{u_3\, x_3}{D})}{u_3} + \frac{1-exp(-\frac{u_4\, x_4}{D})}{u_4})})^{-1} \\ v_{3/4} &= \beta(1 + \frac{c}{D}\frac{exp(-\frac{u_{3/4}x_{3/4}}{D})exp(-\frac{u_m\, d}{D})}{\frac{1-exp(-\frac{u_m\, d}{D})}{u_m} + \frac{1-exp(-\frac{u_1\, x_1}{D})}{u_1} + \frac{1-exp(-\frac{u_2\, x_2}{D})}{u_2} + exp(-\frac{u_m\, d}{D})\,(\frac{1-exp(-\frac{u_3\, x_3}{D})}{u_3} + \frac{1-exp(-\frac{u_4\, x_4}{D})}{u_4})})^{-1} \end{aligned} \tag{S-14}$$

where we have rescaled $c = \frac{c_{tot}}{c_s}$, and the actin flow speeds $u_i$ are given by Eq.7 and Eq.8.

### S-2. Critical local actin polymerization activity for migration

#### Critical local actin polymerization activity for symmetry breaking

As in our previous model for cells on one junction, for cells that are spanning symmetrically on more junctions, there are also critical values of $\beta$ which decide whether the cell can break the symmetry and start migrating. The $\beta_c$ is determined by two lengths of the cell: the critical length of polarization due to the force balance, $L_p$, and the critical length of polarization due to the redistribution of the polarity cue, $L_c$.

$L_p$ is calculated by the balance between the protrusion force and the elasticity of the cell. The cell has a tendency to elongate when its length is shorter than $L_p$. For the cases where cells occupy multiple nodes, the expression of $L_p$ is the same as when cells migrate on a one-dimensional line, and when cells migrate on a single junction

$$L_p = \frac{1}{2}(1-c) + \frac{\beta}{2k} + \sqrt{c + (\frac{1-c}{2} + \frac{\beta}{2k})^2} \tag{S-15}$$

On the other hand, when the cell length is longer than $L_c$, the actin treadmilling flows of arms will deviate from the uniform solutions. In the instance of this symmetry breaking event, all the arms have the equal length, but there will be a small bias in some of the arms. We can derive $L_c$ based on this event, and then the $\beta_c$ is obtained by equating $L_p$ and $L_c$.

For cells migrating on a one-dimensional line, and for cells on a single junction, the $\beta_c$ is given by [3, 4]

$$\beta_c(n_j = 0) = \frac{D}{2c} + ck + \frac{ck^2}{2D} + \frac{D-ck}{2cD}\sqrt{D^2 + 2c(1+2c)Dk + c^2k^2} \tag{S-16}$$

$$\beta_c(n_j=1)=\frac{D}{c}+ck+\frac{ck^2}{4D}+\frac{2D-ck}{4cD}\sqrt{4D^2+4c(1+2c)Dk+c^2k^2} \tag{S-17}$$

Next, we will derive the $\beta_c$ in the 2-junction case as an example. We assume that $x_i=l$, $u_{1/2}=\epsilon$ and $u_{3/4}=-\epsilon$ (Fig. S-1A), then according to Eq.S-1 and Eq.S-2, we have $u_m=u_3+u_4=-2\epsilon$. By substituting these values to Eq.S-14, we obtain the actin treadmilling flows equations

$$\begin{aligned} v_{1/2}&=\beta\Big(1+\frac{c}{D}\frac{exp(-\frac{\epsilon l}{D})}{\frac{1}{2\epsilon}[1-exp(-\frac{2\epsilon d}{D})]+\frac{2}{\epsilon}[1-exp(-\frac{\epsilon l}{D})]+exp(\frac{2\epsilon d}{D})\frac{2}{\epsilon}[1-exp(\frac{\epsilon l}{D})]}\Big)^{-1}\\ v_{3/4}&=\beta\Big(1+\frac{c}{D}\frac{exp(\frac{2\epsilon d}{D})exp(\frac{\epsilon l}{D})}{\frac{1}{2\epsilon}[1-exp(-\frac{2\epsilon d}{D})]+\frac{2}{\epsilon}[1-exp(-\frac{\epsilon l}{D})]+exp(\frac{2\epsilon d}{D})\frac{2}{\epsilon}[1-exp(\frac{\epsilon l}{D})]}\Big)^{-1} \end{aligned} \tag{S-18}$$

Then we can write the global flow of arm 1 (Eq.S-1) as

$$\begin{aligned} \epsilon=&\frac{\beta}{2}\Big(1+\frac{c}{D}\frac{exp(-\frac{\epsilon l}{D})}{\frac{1}{2\epsilon}[1-exp(-\frac{2\epsilon d}{D})]+\frac{2}{\epsilon}[1-exp(-\frac{\epsilon l}{D})]+exp(\frac{2\epsilon d}{D})\frac{2}{\epsilon}[1-exp(\frac{\epsilon l}{D})]}\Big)^{-1}\\ &-\frac{\beta}{2}\Big(1+\frac{c}{D}\frac{exp(\frac{2\epsilon d}{D})exp(\frac{\epsilon l}{D})}{\frac{1}{2\epsilon}[1-exp(-\frac{2\epsilon d}{D})]+\frac{2}{\epsilon}[1-exp(-\frac{\epsilon l}{D})]+exp(\frac{2\epsilon d}{D})\frac{2}{\epsilon}[1-exp(\frac{\epsilon l}{D})]}\Big)^{-1} \end{aligned} \tag{S-19}$$

Expand it in the first order in $\epsilon$

$$\epsilon=\frac{\beta c}{D}\frac{(d+l)(d+4l)}{(c+d+4l)^2}\epsilon+O(\epsilon^2) \tag{S-20}$$

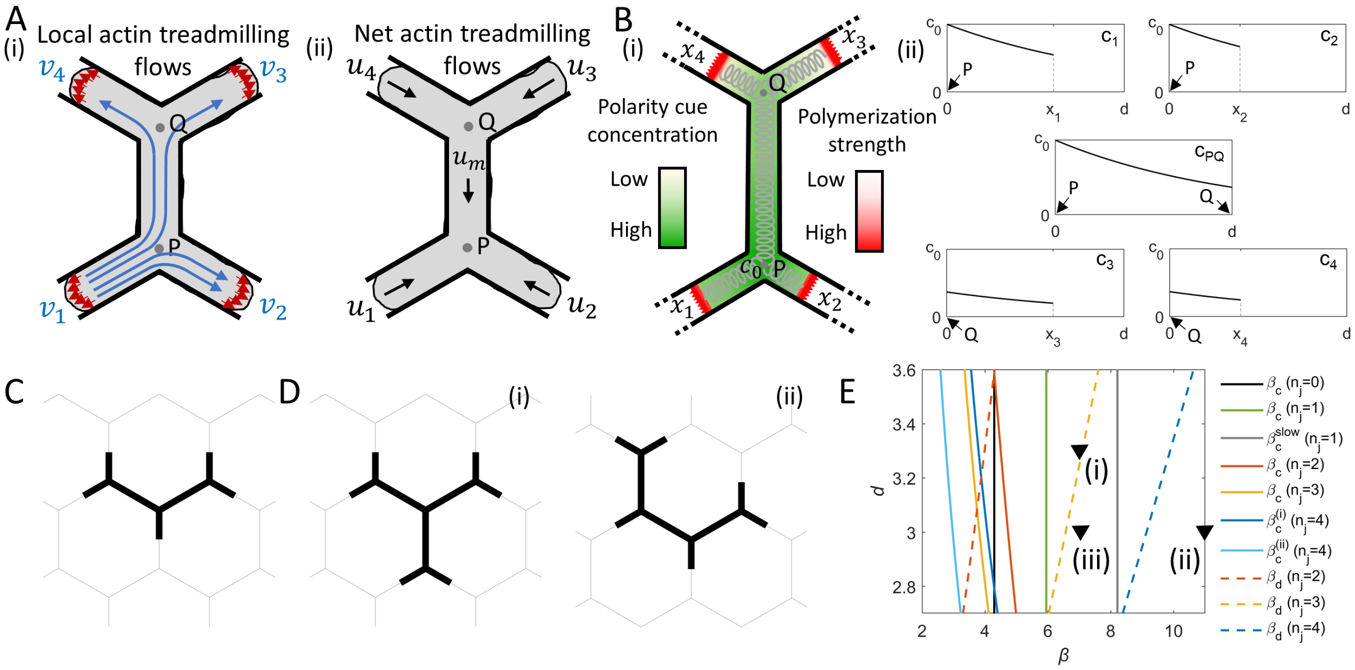

Fig. S-1: (A) (i) Illustrations of the local actin treadmilling flows at each edge of the arms, and the split of the local actin flow of arm 1. (ii) Illustrations of the net actin treadmilling flows in each segment of the cell. (B) (i) An example of the concentration field of the polarity cue, which is affected by the advection flows. The cell elasticity is denoted by the grey springs. (ii) Polarity cue concentration profiles of all the segments in (i), where $d$ is the length of the hexagonal network segments. Node P is the reference point where the polarity cue concentration is fixed to be $c_0$ (Eq.S-12), and node Q is the other junction node that the cell is spanning. C) The cell shape for cells spanning 3 junctions. D) Two possible cell shapes for cells spanning 4 junctions. E) Critical $\beta$s for symmetry breaking and for occupying multiple junctions (Eqs.S-16, S-17, S-23, S-24, S-25, S-26, S-28). Black inverted triangles corresponds to those in Fig. 2A. Parameters: $c=3.85, D=3.85, k=0.8$.

By solving it for $l$, we obtain the solution of the critical length for each arm

$$l_c = \frac{8dD + c(8D - 5d\beta + \sqrt{\beta(16cD - 48dD + 9d^2\beta)}}{8(\beta c - 4D)} \tag{S-21}$$

The critical length of the cell is

$$L_c = 4l_c + d = \frac{8dD + c(8D - 5d\beta + \sqrt{\beta(16cD - 48dD + 9d^2\beta)}}{2(\beta c - 4D)} + d \tag{S-22}$$

By equating $L_p$ (Eq.S-15) and $L_c$ (Eq.S-22), we obtain the $\beta_c$ for symmetry breaking of cells occupying 2 junctions,

$$\begin{aligned}\beta_c(n_j = 2) =& \frac{1}{2c(4D + 3dk)}[16D^2 + k^2c(c - 3d)(3d + 1) + 4Dk(2c^2 + 3d - 3cd) \\ &+ (4D - kc - 3kd)\sqrt{16D^2 + 8c(2c - 3d + 1)Dk + c^2(3d + 1)^2k^2}]\end{aligned} \tag{S-23}$$

By similar derivations, we obtain the $\beta_c$ for the 3-junction case and for the 4-junction case (Fig.S-1C,D),

$$\begin{aligned}\beta_c(n_j = 3) =& \frac{1}{24c(5D + 6dk)}[400D^2 + 9k^2c(c - 8d)(8d + 1) + 120Dk(c^2 + 4d - 4cd) \\ &+ (20D - 3kc - 24kd)\sqrt{400D^2 + 120c(2c - 8d + 1)Dk + 9c^2(8d + 1)^2k^2}]\end{aligned} \tag{S-24}$$

$$\begin{aligned}\beta_{c4}^{(i)}(n_j = 4) =& \frac{1}{8c(8D + 9dk)}[64D^2 + k^2c(c - 9d)(9d + 1) + 8Dk(2c^2 + 9d - 9cd) \\ &+ (8D - kc - 9kd)\sqrt{64D^2 + 16c(2c - 9d + 1)Dk + c^2(9d + 1)^2k^2}]\end{aligned} \tag{S-25}$$

$$\begin{aligned}\beta_{c4}^{(ii)}(n_j = 4) =& \frac{1}{6c(4D + 7dk)}[144D^2 + k^2c(c - 21d)(21d + 1) + 12Dk(2c^2 + 21d - 21cd) \\ &+ (12D - kc - 21kd)\sqrt{144D^2 + 24c(2c - 21d + 1)Dk + c^2(21d + 1)^2k^2}]\end{aligned} \tag{S-26}$$

#### Critical local actin polymerization activity for occupying multiple junctions

To obtain the critical $\beta$ for cells to migrate when occupying multiple junctions, it is also necessary to consider if $\beta$ is large enough to enable cells to achieve the minimum length to occupy a certain number of junctions. For a cell with $N$ ($N \geq 3$) arms, i.e., spanning across $N - 2$ junctions, the minimum length is

$$L_{min} = (N - 3)d \tag{S-27}$$

which is the total length of the segments between the adjacent junctions that the cell is occupying.

By equating the polarization length, $L_p$ (Eq.S-15), and $L_{min}$ (Eq.S-27), we obtain the critical $\beta$ for cells to occupy $N - 2$ junctions

$$\beta_d(n_j = N - 2) = \frac{k[(N - 3)d - 1][(N - 3)d + c]}{(N - 3)d} \tag{S-28}$$

In Fig.S-1E we plot $\beta_c$ and $\beta_d$ for different number of junctions, as well as the critical beta of the slow process in the one-junction case [4], for the range of grid size $d$ that we used in this study. Note that the larger one between $\beta_c$ and $\beta_d$ is the critical $\beta$ for cells to migrate when occupying multiple junctions. For the 2-junction case, the critical $\beta$ is $\beta_c$, while for the 3-junction and 4-junction cases, the critical $\beta$ is $\beta_d$, within the range of $d$ values shown.

#### S-3. Mean-squared-displacement

The mean-squared-displacement (MSD) of the center-of-mass of cells is given by

$$MSD(t) = \langle x(t)^2 + y(t)^2 \rangle = \frac{1}{N_c}\sum_{i=1}^{N_c}(x_i(t)^2 + y_i(t)^2) \tag{S-29}$$

where $N_c$ is the number of cells that considered as an ensemble to be averaged (i.e., we run $N_c$ repetitions of simulations), and $x/y(t)$ is the $x/y$ coordinate of the centroid of ith cell ($i = 1, 2, ..., N_c$) at time t in the simulation. Here we set $N_c = 1000$.

#### S-4. Junction residency time for a specific number of junctions

We show the $\beta - \sigma$ phase diagrams of the average junction residency time for $d = 3.0$ and $d = 3.5$ in the main text. The corresponding phase diagrams of the average junction residency time for a specific number of junctions are shown in Fig.S-2A,B, respectively.

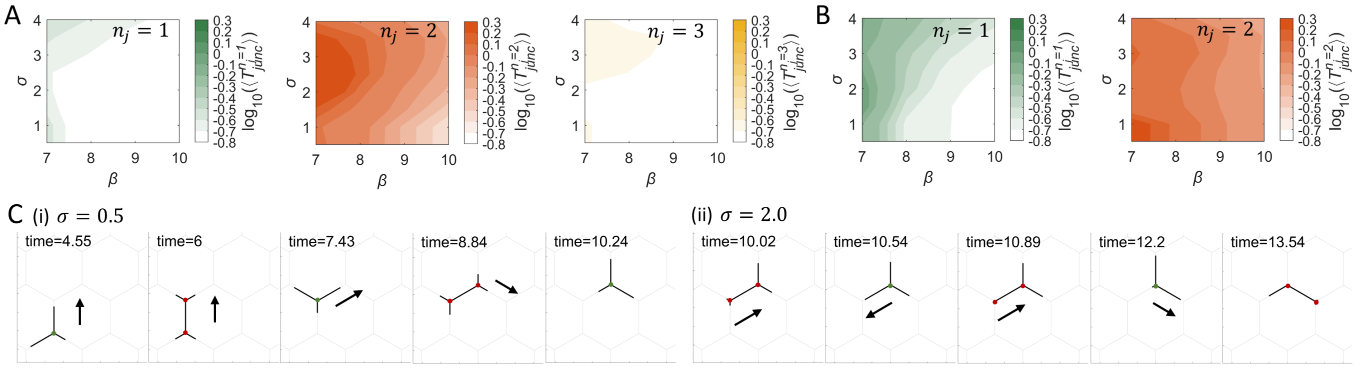


Fig. S-2: The $\beta - \sigma$ phase diagrams of the junction residency time for a specific number of junctions (in log scale). (A) $d = 3.0$. (B) $d = 3.5$. Other parameters: $c = 3.85, D = 3.85, k = 0.8, f_s = 5, r = 5, \kappa = 20, \delta = 250$. (C) Simulation Snapshots, (i) and (ii) for the time stamps (gray dashed lines) in the right panel of Fig. 3E(ii) and (iii), respectively.

#### S-5. Hexagonal residency time

The hexagonal residency time, $T_{hex}$, is defined as the time from the cell's center-of-mass entering a hexagonal grid to leaving it and entering a new one. We show the phase diagram of the average hexagonal residency time, $\langle T_{hex}\rangle$ (Fig.S-3A). Consistent with the trend of $\langle T_{junc}\rangle$ (Fig. 3), the increase of $\beta$ decreases $\langle T_{hex}\rangle$.

We notice that for cells with large $\beta$ and small $\sigma$, the distribution of $T_{hex}$ always have some peaks, which are correlated to some periodic patterns of cell length and number of junctions during the migration process. For example, the $T_{hex}$ distribution for $(\beta, d, \sigma) = (10.0, 2.5, 0.5)$ has two discrete peaks (Blue/red in Fig.S-3B(i)), corresponding to the first and second half of the circulation of cell length and number of junctions (blue and red lines in Fig.S-3B(ii)). In Fig.S-3C, we show the snapshots for the shape and the center-of-mass of the cell in a circulation. The center-of-mass is marked by the red dot, and the grid that it is occupying is colored in blue/red, which corresponds to the blue/red peak in Fig.S-3B(i). Note that we ignore the small peak near $T_{hex} \approx 0$ (Gray peak in Fig.S-3B(ii)), which not indicates to periodic patterns. They originate from the small fluctuation of arm lengths when the cell shape is nearly symmetric and the center-of-mass is close to the border of two adjacent grid. In this case, the slight drift of the center-of-mass may lead to the stop of the residency on a grid.

The peaks of $T_{hex}$ distributions become less significant as $\beta$ decreases, because there are almost no oscillations in the cell length. An example for $(\beta, d, \sigma) = (7.0, 2.5, 0.5)$ is shown in Fig.S-3D(i). The peaks shift left and connect with each other when $\sigma$ is large, because the noise makes the center-of-mass drifts more irregularly. An example for $(\beta, d, \sigma) = (10.0, 2.5, 2.0)$ is shown in Fig.S-3D(ii).

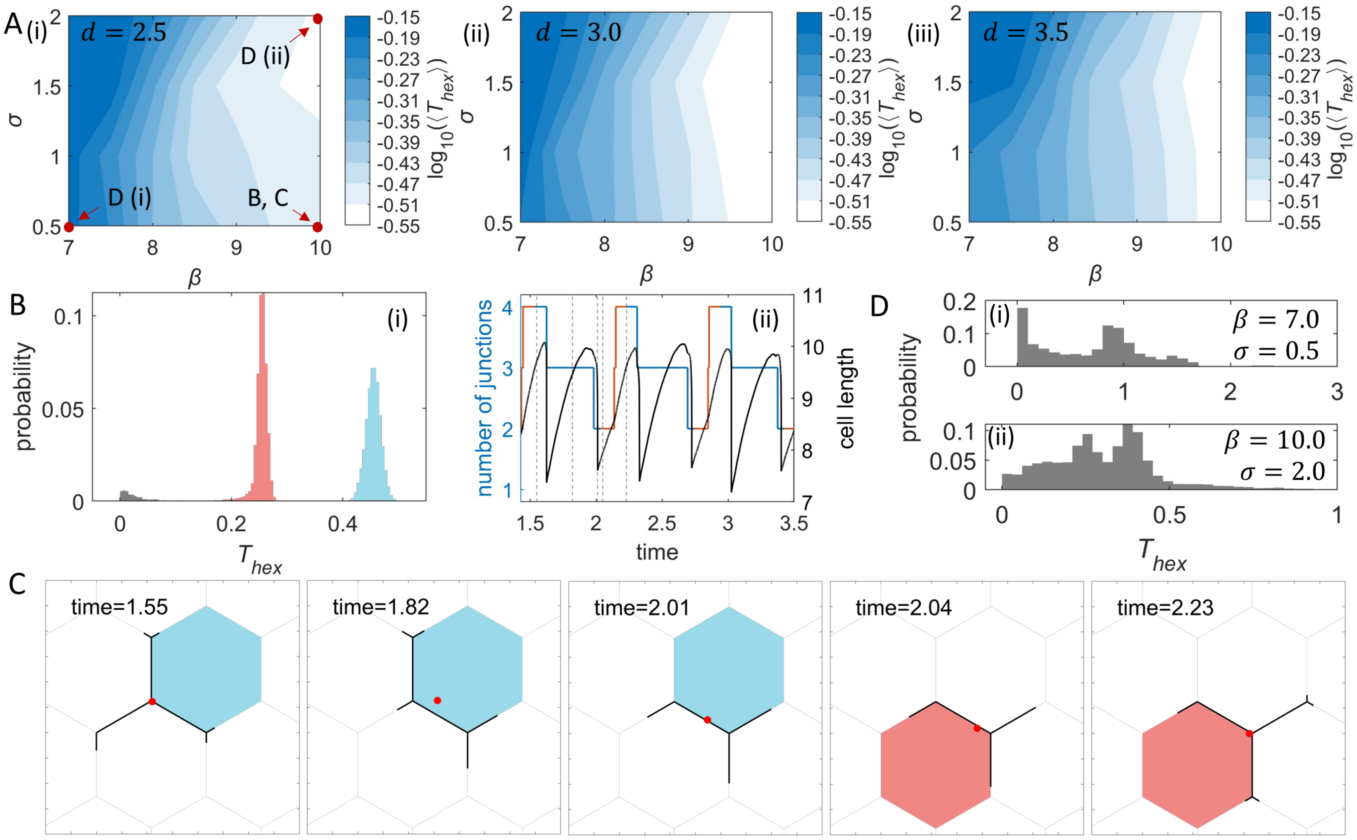


Fig. S-3: Analysis of the hexagonal residency time. A) The $\beta - \sigma$ phase diagram of $\langle T_{hex} \rangle$ (in log scale) for different sizes of grids. (i), (ii) and (iii) correspond to the grid size $d = 2.5$, $d = 3.0$ and $d = 3.5$. B) Analysis of the peaks of $T_{hex}$ distribution for $(\beta, d, \sigma) = (10.0, 2.5, 0.5)$. i) The $T_{hex}$ distribution. ii) Time series of the number of arms and cell length in a simulation. Blue/red lines of the number of arms correspond to the blue/red peaks in (i). C) Snapshots of cell shape in a simulation, (i-v) corresponding to the five time stamps (gray dashed lines) in (ii) respectively. The red point indicates the centroid of the cell. The colored grid is the grid that the centroid is locating in, and blue/red correspond to the blue/red peaks i of $T_{hex}$ in B(i). D) The $T_{hex}$ distribution for $d = 2.5$. (i) and (ii) correspond to $(\beta, \sigma) = (7.0, 0.5)$ and $(\beta, \sigma) = (10.0, 2.0)$ respectively. Parameters: $c = 3.85, D = 3.85, k = 0.8, f_s = 5, r = 5, \kappa = 20, \delta = 250$.

### S-6. Micropatterns of HUVEC and macrophages

The micropatterns of HUVEC and macrophages in our experiments has been performed as described the "Experimental Methods" section of the main text, and shown schematically in Fig.S-4. Note that we have adapted the size of the hexagonal array to allow the HUVEC and macrophages to span simultaneously several junctions. Because macrophages are smaller than HUVEC we have reduced the size of the hexagons previously used [4], as described in Fig.S-4.

### Supplementary movies

Movie S1: Macrophages migrating *in vivo* in the basal layer of keratinocytes in the zebrafish tailfin.
Movie S2: Fluorescent movie of a macrophage migrating *in vivo*.
Movie S3: Simulation of the migration of the same cell on grids with different sizes (Fig. 2B(i),(iii)).
Movie S4: Experiment of a migrating HUVEC (Fig. 4A).
Movie S5: Simulation of the migrating HUVEC in Fig. 4A (Fig. 4B).
Movie S6: Experiment of a migrating macrophage (Fig. 4E).
Movie S7: Simulation of the migrating macrophage in Fig. 4E (Fig. 4F).
Movie S8: Experiment of a migrating HUVEC treated by the inhibitor of calpain (Fig. 5A).
Movie S9: Simulation of the migrating HUVEC in Fig. 5A (Fig. 5B).

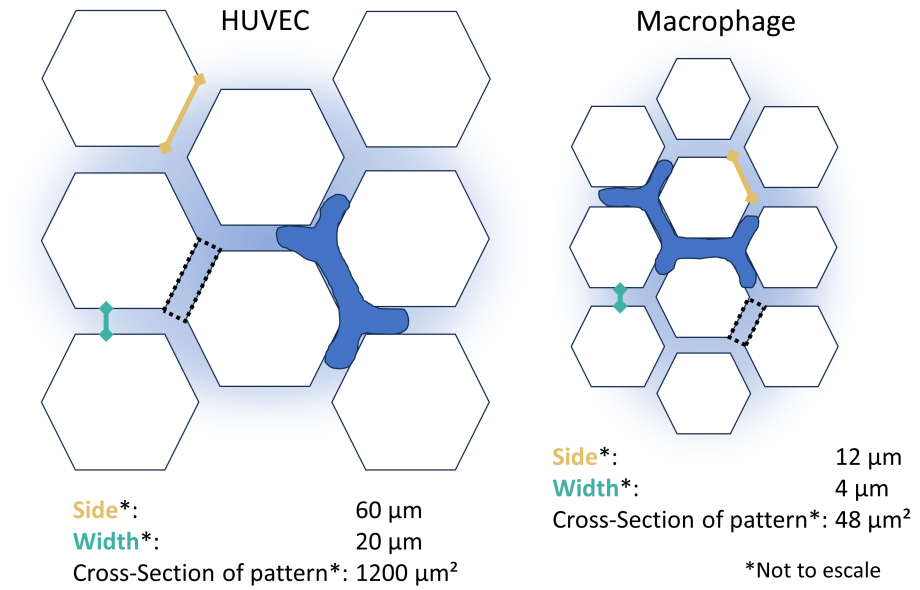


Fig. S-4: Micropatterns of HUVEC and macrophages. Left: Hexagonal micropattern of the HUVEC. Right: Hexagonal micropattern of the macrophages. Note the differences in the size of the hexagonal array for each cell type.

Movie S10: Experiment of a migrating HUVEC (Fig. S-5A).
Movie S11: Simulation of the migrating HUVEC in Fig. S-5A (Fig. S-5B).
Movie S12: Experiment of a migrating HUVEC (Fig. S-5C).
Movie S13: Simulation of the migrating HUVEC in Fig. S-5C (Fig. S-5D).
Movie S14: Experiment of a migrating macrophage (Fig. S-5E).
Movie S15: Simulation of the migrating macrophage in Fig. S-5E (Fig. S-5F).
Movie S16: Experiment of a weakly motile HUVEC (Fig. S-6A).
Movie S17: Simulation of the weakly motile HUVEC in Fig. S-6A (Fig. S-6B).
Movie S18: Experiment of a weakly motile HUVEC (Fig. S-6C).
Movie S19: Simulation of the weakly motile HUVEC in Fig. S-6C (Fig. S-6D).
Movie S20: Experiment of a weakly motile HUVEC (Fig. S-6E).
Movie S21: Simulation of the weakly motile HUVEC in Fig. S-6E (Fig. S-6F).
Movie S22: Experiment of a migrating HUVEC treated by the inhibitor of calpain (Fig. S-7A).
Movie S23: Simulation of the migrating HUVEC in Fig. S-7A (Fig. S-7B).

[1] P. Maiuri, J.-F. Rupprecht, S. Wieser, V. Ruprecht, O. Bénichou, N. Carpi, M. Coppey, S. De Beco, N. Gov, C.-P. Heisenberg, et al., Cell **161**, 374 (2015).
[2] I. Lavi, M. Piel, A.-M. Lennon-Duménil, R. Voituriez, and N. S. Gov, Nature Physics **12**, 1146 (2016).
[3] J. E. Ron, P. Monzo, N. C. Gauthier, R. Voituriez, and N. S. Gov, Physical Review Research **2**, 033237 (2020).
[4] J. E. Ron, M. Crestani, J. M. Kux, J. Liu, N. Al-Dam, P. Monzo, N. C. Gauthier, P. J. Sáez, and N. S. Gov, Nature Physics pp. 1–11 (2024).

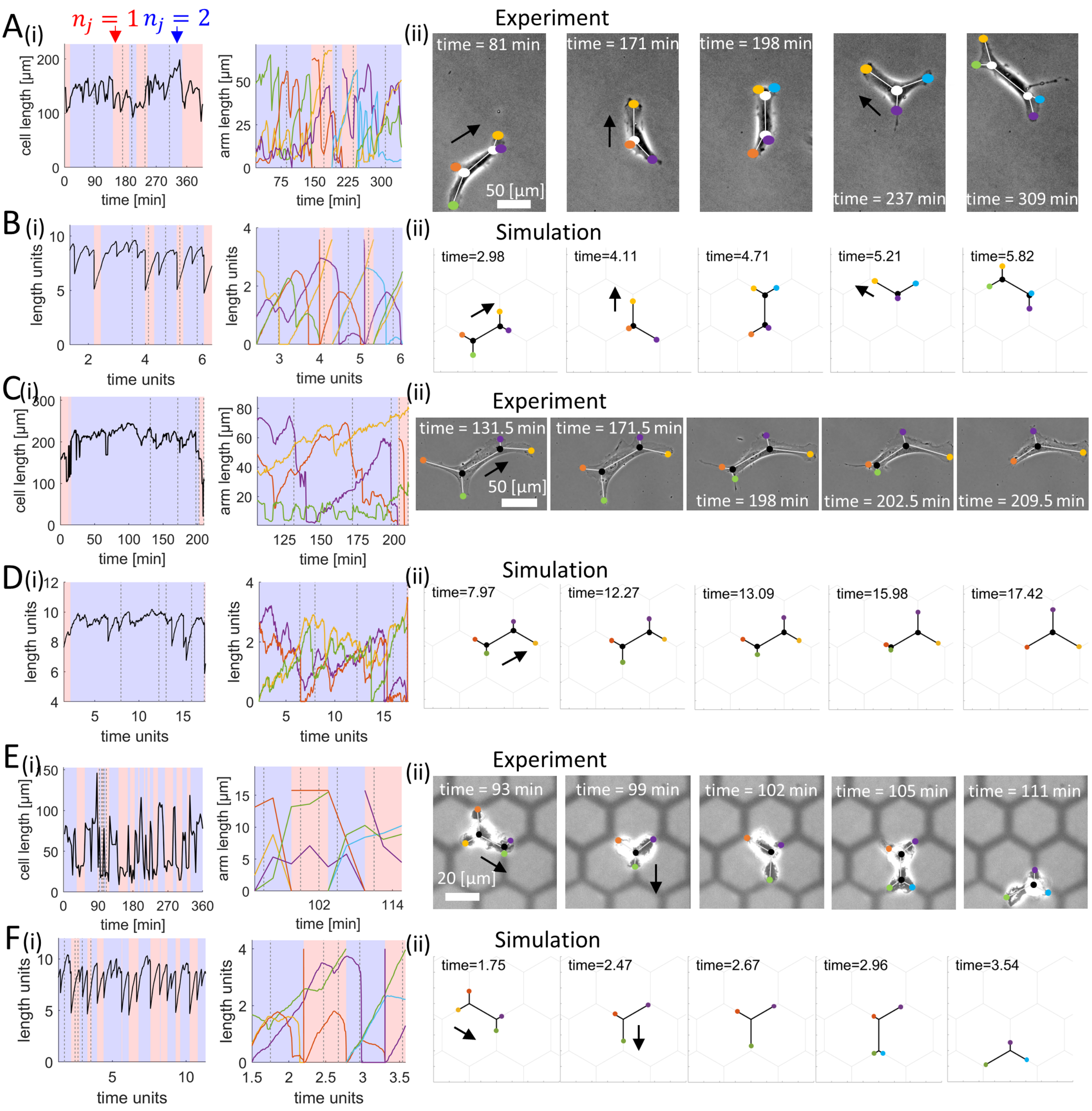


Fig. S-5: Comparisons between cell shapes and migration patterns in simulations and experiments on HUVEC and macrophages. (A,C) Experiments on migrating HUVEC (A, Movie S10; C, Movie S12). Hexagon side: 60 μm. Inter-hexagon width: 20 μm. (B,D) Simulations of non-migrating HUVEC (B, Movie S11; D, Movie S13). i) Dynamics of the total cell length and the arm lengths. ii) Snapshots of the cell, corresponding to gray dashed lines in (i). Key simulation parameters: $(\beta, d, \sigma, k) = (9.0, 3.6, 1.8, 0.8)$ in (B), and $(\beta, d, \sigma, k) = (7.0, 3.5, 2.7, 0.7)$ in (D). Other simulation parameters: $c = 3.85, D = 3.85, f_s = 5, r = 5, \kappa = 20, \delta = 250$. (E) Experiment on a migrating macrophage (Movie S14). Hexagon side: 12 μm. Inter-hexagon width: 4 μm. (F) Simulations of the migrating macrophage (Movie S15). i) Dynamics of the total cell length and the arm lengths. ii) Snapshots of the cell, corresponding to gray dashed lines in (i). Key simulation parameters: $(\beta, d, \sigma) = (10.0, 4.0, 2.5)$. Other simulation parameters: $c = 3.85, D = 3.85, k = 0.8, f_s = 5, r = 5, \kappa = 20, \delta = 250$.

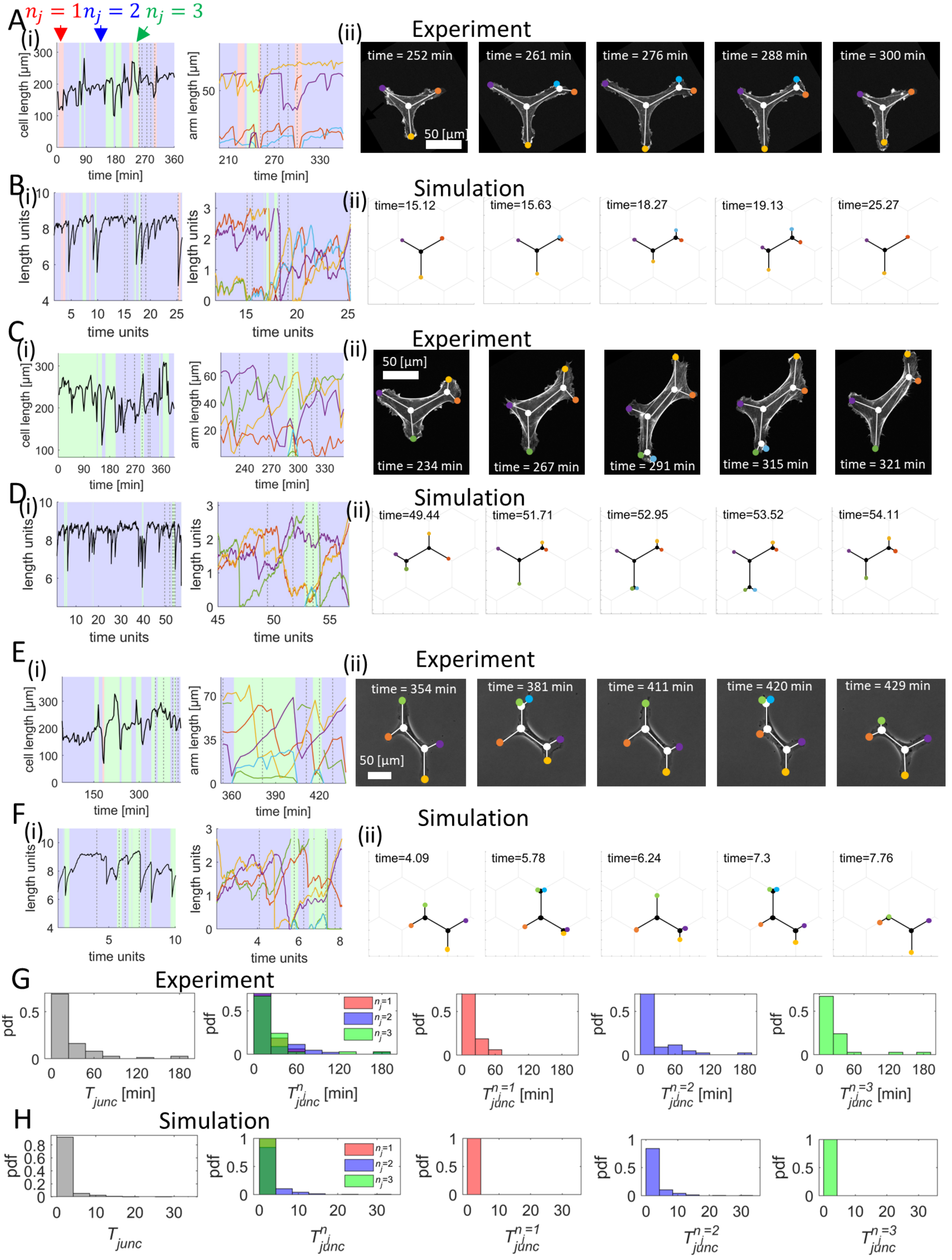

Fig. S-6: Comparisons between cell shapes dynamics in simulations and experiments for weakly motile HUVEC. (A,C,E) Experiments on non-migrating HUVEC (A, Movie S16; C, Movie S18; E, Movie S20). Hexagon side: 60 µm. Inter-hexagon width: 20 µm. (B,D,F) Simulations of non-migrating HUVEC (B, Movie S17; D, Movie S19; F, Movie S21). i) Dynamics of the total cell length and the arm lengths. ii) Snapshots of the cell, corresponding to gray dashed lines in (i). Key simulation parameters: $(\beta, d, \sigma) = (7.0, 3.0, 1.8)$ in (B), $(\beta, d, \sigma) = (7.0, 2.7, 2.1)$ in (D), and $(\beta, d, \sigma) = (7.6, 2.7, 2.0)$ in (F). Other simulation parameters: $c = 3.85, D = 3.85, k = 0.8, f_s = 5, r = 5, \kappa = 20, \delta = 250$. (G) Distribution (probability density function, pdf) of $T_{junc}$ (leftmost) and distribution of $T_{junc}^{n_j}$ (right ones) in the experimental regime in which the cells spans 1, 2 or 3 junctions. A total of 3 cells across 1725 minutes were used to obtain these distributions. (H) Distribution (pdf) of $T_{junc}$ (leftmost) and distribution of $T_{junc}^{n_j}$ (right ones) from a long simulation with parameters of (B).

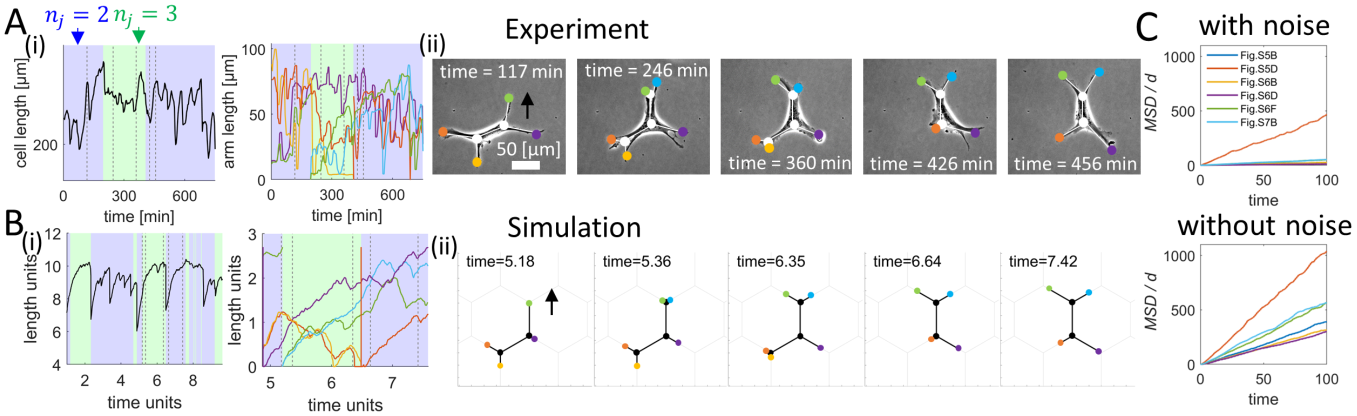


Fig. S-7: Comparisons between cell shapes dynamics in simulations and experiments for HUVEC with larger cell-substrate adhesiveness. (A) Experiment on HUVEC treated by the inhibitor of calpain (Movie S22). (B) Simulation of HUVEC with larger cell-substrate adhesiveness (Movie S23). Hexagon side: 60 μm. Inter-hexagon width: 20 μm. i) Dynamics of the total cell length and the arm lengths. ii) Snapshots of the cell, corresponding to gray dashed lines in (i). Key simulation parameters: $(\beta, d, \sigma, r) = (8.5, 2.7, 2.5, 7.0)$. Other simulation parameters: $c = 3.85, D = 3.85, k = 0.8, f_s = 5, \kappa = 20, \delta = 250$. (C) Comparisons of $MSD$ of the simulated HUVEC of Fig. S-5, Fig. S-6 and Fig. S-7, with the noise levels identical to those used in the corresponding simulations (top) or without the noise (bottom).
